# Blimp-1 and c-Maf regulate common and unique gene networks to protect against distinct pathways of pathobiont-induced colitis

**DOI:** 10.1101/2023.09.14.557705

**Authors:** Marisol Alvarez-Martinez, Luke S. Cox, Claire F. Pearson, William J. Branchett, Probir Chakravarty, Xuemei Wu, Hubert Slawinski, Alaa Al-Dibouni, Vasileios A. Samelis, Leona Gabryšová, Simon L. Priestnall, Alejandro Suárez-Bonnet, Anna Mikolajczak, James Briscoe, Fiona Powrie, Anne O’Garra

**Affiliations:** Immunoregulation and Infection Laboratory, The Francis Crick Institute, London, UK; Kennedy Institute of Rheumatology, University of Oxford, Oxford, UK; Computational Biology Laboratory, The Francis Crick Institute, London, UK; Advanced Sequencing Facility, The Francis Crick Institute, London, UK; Department of Pathobiology and Population Sciences, Royal Veterinary College, London, UK; 6Experimental Histopathology, The Francis Crick Institute, London, UK; Developmental Dynamics Laboratory, The Francis Crick Institute, London, UK; National Heart and Lung Institute, Imperial College London, London, UK

**Author notes:** Correspondence and requests for materials should be addressed to AOG. Both these authors contributed equally (Marisol Alvarez-Martinez, Luke S. Cox). Both these authors contributed equally (Claire F. Pearson, William J. Branchett).

## Abstract

Intestinal immune responses to commensals and pathogens are controlled by IL-10 to avoid intestinal immune pathology. We show that the transcription factors Blimp-1 *(Prdm-1)* and c-Maf are co-dominant regulators of *Il10* in Foxp3^+^ regulatory T cells, but also negatively regulate proinflammatory cytokines in effector T cells. Mice with T cell-specific deletion of *Prdm-1, Maf* or the combination of both transcription factors did not develop inflammatory intestinal pathologies at the steady state. Double deficient *Prdm1*^fl/fl^*Maf*^fl/fl^*Cd4*^Cre^ mice infected with *Helicobacter hepaticus* developed severe colitis with a major increase in TH1/NK/ILC1 effector genes in lamina propria leucocytes (LPLs), while *Prdm1*^fl/fl^*Cd4*^Cre^ and *Maf*^fl/fl^*Cd4*^Cre^ mice showed mild/moderate pathology and a less-marked Type I effector response. LPLs from infected *Maf*^fl/fl^*Cd4*^Cre^ mice showed increased *Il17a* expression and an accompanying increase in granulocytes and myeloid cells, which was less marked in *Prdm1*^fl/fl^*Maf*^fl/fl^*Cd4*^Cre^ mice, with increased T cell-myeloid-neutrophil interactions inferred from scRNA-seq analysis and confirmed by immunofluorescent analysis of colon sections. Genes over-expressed in human IBD showed differential expression in the LPL from infected mice in the absence of *Prdm1* or *Maf,* revealing potential pathobiologic mechanisms of human disease.

---

The immune response has evolved to protect the host against infection however, regulatory mechanisms are in place to prevent untoward inflammation and host damage. One such mechanism is provided by the cytokine IL-10, which has predominantly anti-inflammatory properties and suppresses immune responses to pathogens and pathobionts^1, 2, 3, 4, 5^. Mice deficient in IL-10 (*Il-10-/-*) can develop colitis^6^. This is less evident in specific-pathogen-free (SPF)-reared *Il-10-/*- mice and not evident in germ-free mice^7^, but is triggered by pathobionts, such as *Helicobacter hepaticus (H. hepaticus)*^8^. T cell-derived IL-10 is critical for control of the immune response with T cell-specific IL-10 mutant mice developing colitis to a similar level as *Il-10-/-* mice^9^. Rare loss-of-function mutations in the *Il10, Il10ra*, or *Il10rb* genes result in inflammatory bowel disease (IBD) in childhood, although it is unclear whether infection by pathobionts contributes to these pathologies^10^. Genome-wide association studies (GWASs) have identified more than 230 loci linked to human IBD, including those associated with pro-inflammatory cytokines and transcription factors upstream of immune effector molecules^11^ and transcription factors such as Blimp-1, encoded by *Prdm1,* where rare missense mutations in PR domain-containing 1 (PRDM1) have been associated with IBD^12^.

Both common and cell-specific transcriptional mechanisms are in place to regulate the expression of *Il10* and proinflammatory gene expression in T cells to ensure a controlled immune response to pathogens and/or other insults^1, 3, 4, 5, 13, 14^ (Cox, Alvarez, O’Garra, Wellcome Open Research, 2023, under review). Since transcription factors have multiple gene targets, it is possible that transcription factors that positively regulate *Il10* simultaneously repress proinflammatory cytokine expression in T cells, thus restraining immune responses to pathogens and pathobionts. For example, the transcription factor c-Maf, encoded by *Maf*, has been shown to induce *Il10* expression directly in multiple T cell subsets both *in vitro* and *in vivo*^1^, whilst also acting as a negative regulator of *Il2*^13^ and other proinflammatory cytokines and described as an immune repressor^15^. However, the deletion of *Maf* in T cells or Treg was not reported to show any signs of spontaneous inflammation^13, 16^, whereas in other studies aged mice with T cell-specific deletion of *Maf* or mice with *Maf* deleted in Tregs showed signs of intestinal inflammation^17,18^. The transcription factors, c-Maf and Blimp-1 are dominant shared coregulators of *Il10* gene expression in multiple T cell subsets^14^(Cox, Alvarez, O’Garra, Wellcome Open Research, 2023, under review). While T cell-specific deletion of *Prdm1* has been reported to result in spontaneous intestinal inflammation and colitis ^19, 20, 21, 22, 23^, other studies reported no intestinal inflammation in these mice ^14, 24, 25^. The spontaneous severe colitis in T cell-specific deletion of *Prdm1* mice has been associated with increased frequencies of Th17 cells^26^. Although, Blimp-1 functions as a molecular switch to prevent inflammatory activity in Foxp3^+^RORgt^+^ regulatory T cells (Treg)^27^, deletion of *Prdm1* in Treg did not result in severe intestinal inflammation and colitis^28^. More recently, spontaneous colitis has been reported in mice with T cell-specific deletion of the combination of both *Prdm1* and *Maf* and this pathology was associated with a unique cluster of Treg cells and the abrogation of *Il10* expression^14^.

We report here that mice with T cell-specific deletion of *Prdm1, Maf* or the combination of both *Prdm1* and *Maf* do not exhibit inflammation of the colon in the steady state. However, upon infection with *H. hepaticus,* an absence of *Prdm1* resulted in mild pathology, with absence of *Maf* resulting in moderate pathology and the absence of both *Prdm1* and *Maf* showing the greatest pathology. We interrogated the immune response underpinning the pathology in the different T cell-specific transcription factor deficient mice in response to *H. hepaticus* infection using bulk tissue RNA sequencing (RNA-Seq) and single cell-RNA-Seq (scRNA-Seq) of lamina propria leucocytes (LPLs), complemented by RNA-Seq and ATAC-Seq analysis of purified CD4^+^ T cells from the LPLs and validated key findings using flow cytometry and immunofluorescence staining of colon tissue. LPLs from the colons of the double deficient *Prdm1*^fl/fl^*Maf*^fl/fl^*Cd4*^Cre^ *H. hepaticus* infected mice, showed a major increase in genes associated with TH1 and NK/ILC1 effector function including IFN-γ and GM-CSF, but this was lower in *Prdm1*^fl/fl^*Cd4*^Cre^ and *Maf*^fl/fl^*Cd4*^Cre^ mice. By contrast, LPLs from *H. hepaticus* infected *Maf*^fl/fl^*Cd4*^Cre^ mice showed increased expression of *Il17a* and a pronounced signature of innate immunity and neutrophils and myeloid cells, which was also present but less marked in the double *Prdm1*^fl/fl^*Maf*^fl/fl^*Cd4*^Cre^ LPLs. Accompanying this response was an increase in myeloid-neutrophil and myeloid-T cell interaction inferred by scRNA-seq data and confirmed using immunofluorescent staining of colon sections. Genes identified as over-expressed in human IBD RNA-Seq datasets were found to be differentially perturbed in the LPLs of mice with T cell-specific deficiencies in either *Prdm1, Maf* or both transcription factors, potentially reflecting different pathobiological mechanisms that are relevant in human IBD.

## RESULTS

### Blimp-1 and c-Maf control colitis through effects on lymphoid and myeloid cells

*Prdm1*^fl/fl^*Cd4*^Cre^, *Maf*^fl/fl^*Cd4*^Cre^ and *Prdm1*^fl/fl^*Maf*^fl/fl^*Cd4*^Cre^ mice do not exhibit any signs of intestinal inflammation in the “steady state” (Extended Fig. 1a). Upon infection with *H. hepaticus* all these T cell-specific transcription factor deficient mice developed colitis with varying degrees of pathology, with double deficient *Prdm1*^fl/fl^*Maf*^fl/fl^*Cd4*^Cre^ mice developing the most severe disease, and the *Prdm1*^fl/fl^*Cd4*^Cre^ and *Maf*^fl/fl^*Cd4*^Cre^ mice each developed mild and moderate colitis respectively (supported by additional experiments, as compared to *Prdm1*^fl/fl^*Maf*^fl/fl^ control mice (hereafter referred to as control mice), which showed no inflammation or colitis (Fig. 1a - c and Extended Data Supplementary Table 1). This was accompanied by a graded increase in LPL numbers and CD4^+^ T cells counts which was most pronounced in the infected double deficient *Prdm1*^fl/fl^*Maf*^fl/fl^*Cd4*^Cre^ mice, but also observed in the infected single *Prdm1*^fl/fl^*Cd4*^Cre^ and *Maf*^fl/fl^*Cd4*^Cre^ mice (Fig. 1d, e).

**Fig. 1.**
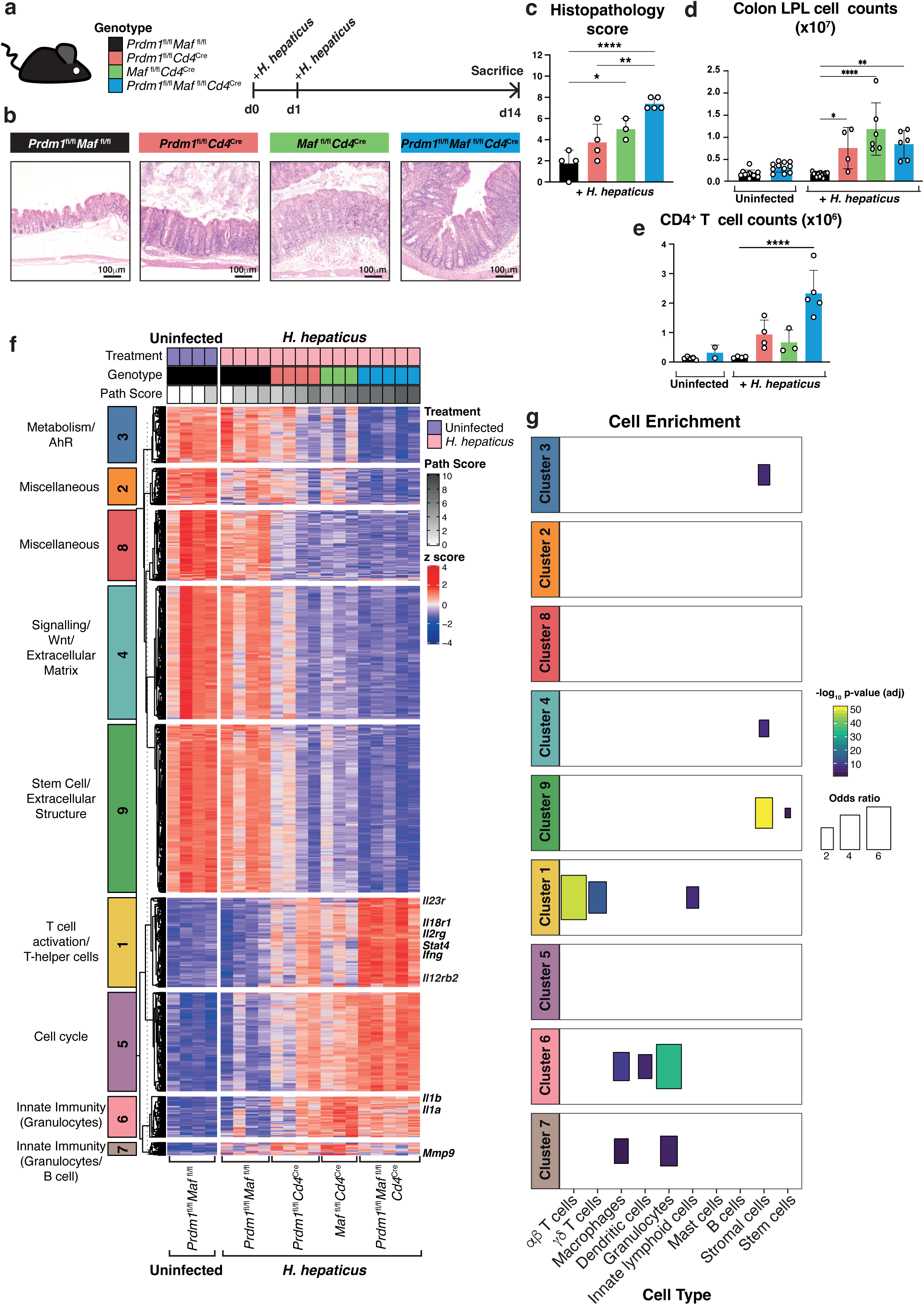
Blimp-1 and c-Maf are required for the protection against *H. hepaticus* induced colitis, with their deletion leading to disruption of gene clusters associated with αβ T cells and granulocytes. **a)** Schematic of experimental method used to infect mice with *H. hepaticus* by oral gavage. **b)** Representative H&E colon sections from each genotype following infection with *H. hepaticus* for 14 days with **c)** the corresponding colon histopathology scores, **d)** colon LPL cell counts and **e)** total CD4+ T cell counts for each group. Each dot within the barplots represents an individual mouse analyzed. Graph shows mean ±s.d., analyzed by one-way ANOVA followed by Tukey’s post-hoc test (*=p value ≤ 0.05, **=p value ≤ 0.01, ***=p value ≤ 0.001, ****=p value ≤ 0.0001). Scale bar = 100μm. **f-g)** Bulk tissue RNA-seq was performed on total colon LPL isolated from uninfected *Prdm1*^fl/fl^*Maf*^fl/fl^ control and *H. hepaticus* infected *Prdm1*^fl/fl^*Maf*^fl/fl^ mice and mice with *Cd4*^Cre^-mediated deletion of either *Prdm1*, *Maf*, or both *Prdm1* and *Maf*. **f)** Heatmap of expression values (represented as z-scores) of differentially expressed genes identified in *H. hepaticus* infected mice compared to uninfected *Prdm1*^fl/fl^*Maf*^fl/fl^ controls (fold change >=1.5 and BH adjusted p value<0.05), partitioned into 9 clusters using *k*-means clustering. Pathology scores associated to each mouse are shown at the top of the heatmap. **g)** Enrichment of cell-type signatures (taken from Singhania *et al*. 2019) was assessed for each of the clusters in **f)** using a Fisher’s exact test. Only statistically significant enriched signatures (BH adjusted p value <0.05) were plotted for visualization. Data from n=3-5 mice.

Since Blimp-1 and c-Maf induce *Il10* gene expression^14^(Cox, Alvarez, O’Garra 2023, Wellcome Open Research, under review) whilst negatively regulating a large network of proinflammatory cytokines (Cox, Alvarez, O’Garra 2023, Wellcome Open Research, under review), it was important to determine whether the effects of T cell-specific deletion of *Prdm1, Maf* or the combination of *Prdm1 and Maf,* resulted in graded increased pathology and inflammation in *H. hepaticus* infected mice due to abrogation of IL-10 signalling. To address this, the T cell-specific transcription factor deficient and control mice were infected with *H. hepaticus* in the presence of anti-IL- 10R blocking antibody (mAb) or isotype matched mAb control (Extended Fig. 1b right hand side). Blockade of IL-10R signalling resulted in moderate to severe pathology in the colons of wild type control mice and resulted in a slight increase in pathology in the single *Prdm1*^fl/fl^*Cd4*^Cre^ and *Maf*^fl/fl^*Cd4*^Cre^ mice infected with *H. hepaticus*. However, the most severe pathology was still observed in the infected double-deficient *Prdm1*^fl/fl^*Maf*^fl/fl^*Cd4*^Cre^ mice in the presence or absence of anti-IL-10R mAb with no significant increase in pathology observed in mice administered anti-IL-10R mAb as compared those given isotype control Mab (Extended Fig.1b, right hand side). These findings suggest that the high level of intestinal pathology observed in the *Prdm1*^fl/fl^*Maf*^fl/fl^*Cd4*^Cre^ mice result from effects of both transcription factors on other immune factors in addition to their co-dominant role in *Il10* gene regulation.

To dissect the mechanisms underlying the pathology observed in the different T cell-specific transcription factor deficient mice, we performed RNA-seq analysis on LPLs isolated from the colons of *H. hepaticus* infected and uninfected mice (Extended Data Fig. 1c, d; Fig. 1f and Extended Data Supplementary Table 2). Compared to uninfected control mice, *H. hepaticus* infected control mice showed only a minor number of 5 differentially expressed genes (5 DEG). Major differences were observed in the *H. hepaticus* infected T cell-specific transcription factor deficient mice, with the number of DEG increasing with the level of pathology: *Prdm1*^fl/fl^*Cd4*^Cre^ (1207 DEG); *Maf*^fl/fl^*Cd4*^Cre^ (1740 DEG) and *Prdm1*^fl/fl^*Maf*^fl/fl^*Cd4*^Cre^ (3392 DEG)(Extended Data Supplementary Table 2). These formed 9 clusters of similarly regulated DEGs (Fig. 1f; Extended Data Supplementary Table 3; and Extended Data Fig. 1e) with the associated pathology scores for each mouse shown at the top of heatmap (Fig. 1f; Extended Data Supplementary Table 1). Gene expression in Clusters 3 (Metabolism/AhR), 4 (Signaling/Wnt/Extracellular Matrix), 9 (Stem Cell/Extracellular Structure) and 2 and 8 (Miscellaneous) were all decreased in the LPL from the *H. hepaticus* infected transcription factor deficient mice as compared to *H. hepaticus* infected and uninfected control mice, accompanying the increasing degree of pathology in the colon (Fig. 1f). Conversely, DEG in Clusters 1 (T cell activation/T-helper cells) and 5 (Cell cycle) were partially increased in LPL from both *Prdm1*^fl/fl^*Cd4*^Cre^ and *Maf*^fl/fl^*Cd4*^Cre^ mice, and further increased in the double deficient *Prdm1*^fl/fl^*Maf*^fl/fl^*Cd4*^Cre^ mice infected with *H. hepaticus,* as compared to the *H. hepaticus* infected and uninfected control mice (Fig. 1f). Differential expression of genes in Clusters 6 (Innate immunity/Granulocytes) and 7 (Innate Immunity/Granulocytes/B cells) was most markedly increased in *H. hepaticus* infected *Maf*^fl/fl^*Cd4*^Cre^ mice and in the double deficient *Prdm1*^fl/fl^*Maf*^fl/fl^*Cd4*^Cre^ mice, albeit to a lesser extent, while barely increased in the infected *Prdm1*^fl/fl^*Cd4*^Cre^ (Fig. 1f; Extended Data Supplementary Tables 3 and 4).

Cell-types associated with each cluster of DEGs in our LPL dataset were identified using the mouse cell-specific RNA-Seq dataset from ImmGen Ultra Low Input (ULI) (GSE109125) as previously described for broad range of infections ^29^. The cell type-specific enrichment data validated the pathway annotation of the clusters representing immune pathways (Fig. 1g and Extended Data Fig1e). Specifically, annotation of DEGs in Cluster 1 by IPA and GO as “T cell activation/T helper cells’, supported by the cell type enrichment data which defined them as αβ- and γ8-T cells and innate lymphoid cells (ILC), the latter sharing many of the genes expressed by the T cells. A dominance of the innate immunity and granulocyte-associated genes identified by pathway annotation tools in Clusters 6 and 7 was also validated showing cell type enrichment for macrophages, dendritic cells and granulocytes (Cluster 6) and macrophages and granulocytes (Cluster 7)(Fig. 1g). Clusters 3, 4 and 9 showed enrichment for stromal cells and additionally Cluster 9 showed a small enrichment for stem cells (Fig. 1g).

To validate these gene signatures and to interrogate in depth the gene expression changes and identify the cellular sources of immune-associated genes induced directly and indirectly by Blimp-1 and c-Maf in discrete cell populations we performed scRNA-Seq on LPL isolated from the colons from an independent *H. hepaticus* infection experiment and uninfected mice (Fig. 2; Extended Fig. 2). Initially, the scRNA-Seq data from the LPL of all groups of infected and uninfected mice were integrated for analysis into a single UMAP revealing 17 distinct cell clusters (Fig. 2a) which were annotated using a combination of the single cell Mouse Cell Atlas^30^, the Immgen data base (GSE109125) and manual curation (Fig. 2a). As with the bulk tissue RNA-Seq data, analysis of the scRNA-Seq data from LPL from uninfected (fl/fl control mice and double deficient *Prdm1*^fl/fl^*Maf*^fl/fl^*Cd4*^Cre^ mice) and infected control *fl/fl* mice, showed distinct profiles to those from infected *Prdm1*^fl/fl^*Cd4*^Cre^, *Maf*^fl/fl^*Cd4*^Cre^ and double deficient *Prdm1*^fl/fl^*Maf*^fl/fl^*Cd4*^Cre^ mice (Fig. 2b; Extended Data Fig. 2a - c). A similar proportion of immune cell types was identified by scRNA-seq in the LPL from the uninfected control mice, uninfected double *Prdm1*^fl/fl^*Maf*^fl/fl^*Cd4*^Cre+^ mice and *H. hepaticus* infected control mice, which was in accordance with no intestinal pathology (Fig. 2b and Extended Data Fig. 2b,c). Moreover, no increase in the numbers of LPL (Fig. 2c) and CD4^+^ T cells (Fig. 2d) assessed by flow cytometry was observed at the steady state, in these mice. However, upon infection with *H. hepaticus,* an increase in intestinal pathology (Fig. 2b) and numbers of LPL and CD4^+^ T cells was observed by flow cytometry (Fig. 2c,d) in *Prdm1*^fl/fl^*Cd4*^Cre^, *Maf*^fl/fl^*Cd4*^Cre^ and double *Prdm1*^fl/fl^*Maf*^fl/fl^*Cd4*^Cre^ as compared to control *fl/fl* mice, again in accordance with the bulk tissue RNA-Seq data (Fig. 1). The scRNA-Seq data revealed a similar increase in the proportion of T cells but additionally revealed greater granularity, particularly showing an increase in a subset of Ctla4 high T cells, in the LPL from *Prdm1*^fl/fl^*Cd4*^Cre^ and *Maf*^fl/fl^*Cd4*^Cre^ mice, with a most marked increase in the infected double deficient *Prdm1*^fl/fl^*Maf*^fl/fl^*Cd4*^Cre^ mice (Fig. 2b). This graded increase was also seen, albeit to a much lesser extent, in proportions of Icos high T cells but not in Cd8 T cells (Fig. 2b). The identity of the annotated cell clusters was further interrogated with the top 10 immune marker genes expressed for selected immune cell clusters (Fig. 2e; Extended Fig. 3a). The NK/ILC cell cluster expressed a distinct discrete set of NK cell-specific genes including *Nkg7, Klrb1b* and the highest level of *Ifng*, confirming their identity as NK/ILC1 cells (Fig. 2e; Extended Fig. 3a). The Ctla4 high and Icos high T cell subsets although clustering separately, broadly shared the expression of the top 10 marker genes (Fig. 2e; Extended Data Fig. 3e). The scRNA-Seq data revealed an increase in granulocytes in *H. hepaticus* infected *Maf*^fl/fl^*Cd4*^Cre^ and double deficient *Prdm1*^fl/fl^*Maf*^fl/fl^*Cd4*^Cre^ only, but not in the infected *Prdm1*^fl/fl^*Cd4*^Cre^ mice or control mice (Fig. 2b), with increased expression of genes associated with neutrophils, including *Acod1, S100a8* and *S100a9* (Fig. 2e; Extended Data Fig. 3; Extended Data Supplementary Table 4). Populations annotated as macrophages and myeloid leukocytes were increased in the LPL from *H. hepaticus* infected *Prdm1*^fl/fl^*Cd4*^Cre^, *Maf*^fl/fl^*Cd4*^Cre^ and double deficient *Prdm1*^fl/fl^*Maf*^fl/fl^*Cd4*^Cre^ mice (Fig. 2b; Extended Data Fig. 2b,c), with increased expression of genes associated with myeloid/innate immune responses such as *Lyz2 (LysM), Csf2rb, Il1r2, Il1b* and *Cd14* (Fig. 2e; Extended Fig. 3a; Extended Data Supplementary Table 4). These scRNA-Seq data thus identified the cellular sources of key gene expression signatures.

**Fig. 2.**
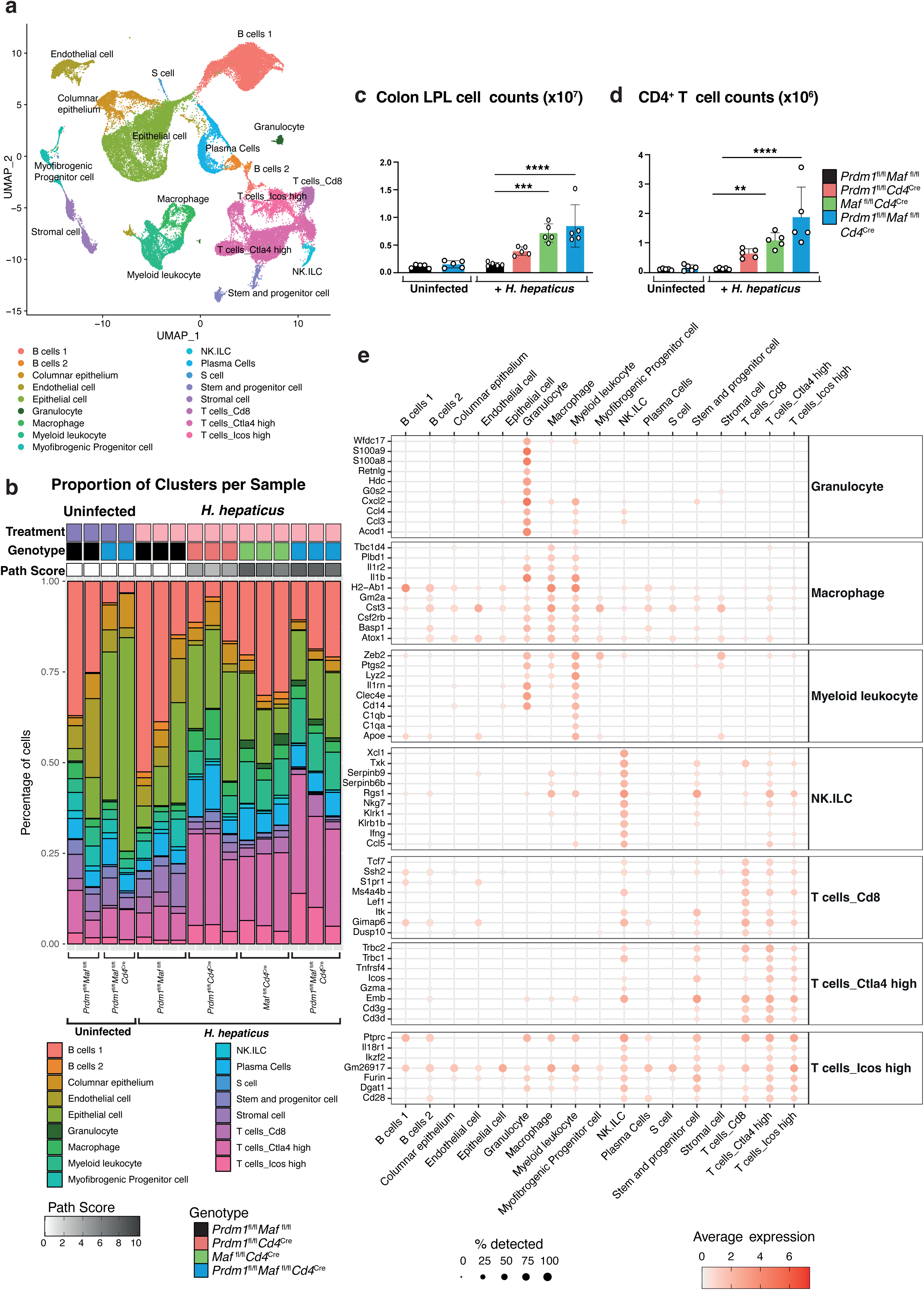
scRNA-seq of total colon LPL from *H. hepaticus* infected mice with *Cd4*^Cre^- mediated deletion of *Prdm1*, *Maf,* or both *Prdm1* and *Maf* reveals qualitative and quantitative differences in cell clusters. **a,b,e)** scRNA-seq was performed on total colon LPL isolated from uninfected *Prdm1*^fl/fl^*Maf*^fl/fl^ and *Prdm1*^fl/fl^*Maf*^fl/fl^*Cd4^C^*^re^ control mice, and *H. hepaticus* infected *Prdm1*^fl/fl^*Maf*^fl/fl^ mice and mice with *Cd4*^Cre^-mediated deletion of either *Prdm1*, *Maf*, or both *Prdm1* and *Maf*. **a)** UMAP visualization of the integrated scRNA-seq from all conditions, colored by the identified/assigned cell cluster. **b)** Barplots representing the proportion of cells in each of the cell clusters per biological replicate within each experimental condition. **c)** Colon LPL cell counts and **d)** total CD4+ T cell counts for each group in the experiment. Each dot within the barplots represents an individual mouse analyzed. Graph shows mean ±s.d., analyzed by one-way ANOVA followed by Tukey’s post-hoc test (*=p value ≤ 0.05, **=p value ≤ 0.01, ***=p value ≤ 0.001, ****=p value ≤ 0.0001). Data from n=5 mice. **e)** Dotplot of the top 10 differentially expressed marker genes of relevant cell clusters from **a)** and colored by the average gene expression across all cell clusters. The dot size represents the percentage of cells per cell cluster expressing the gene in question.

**Fig. 3.**
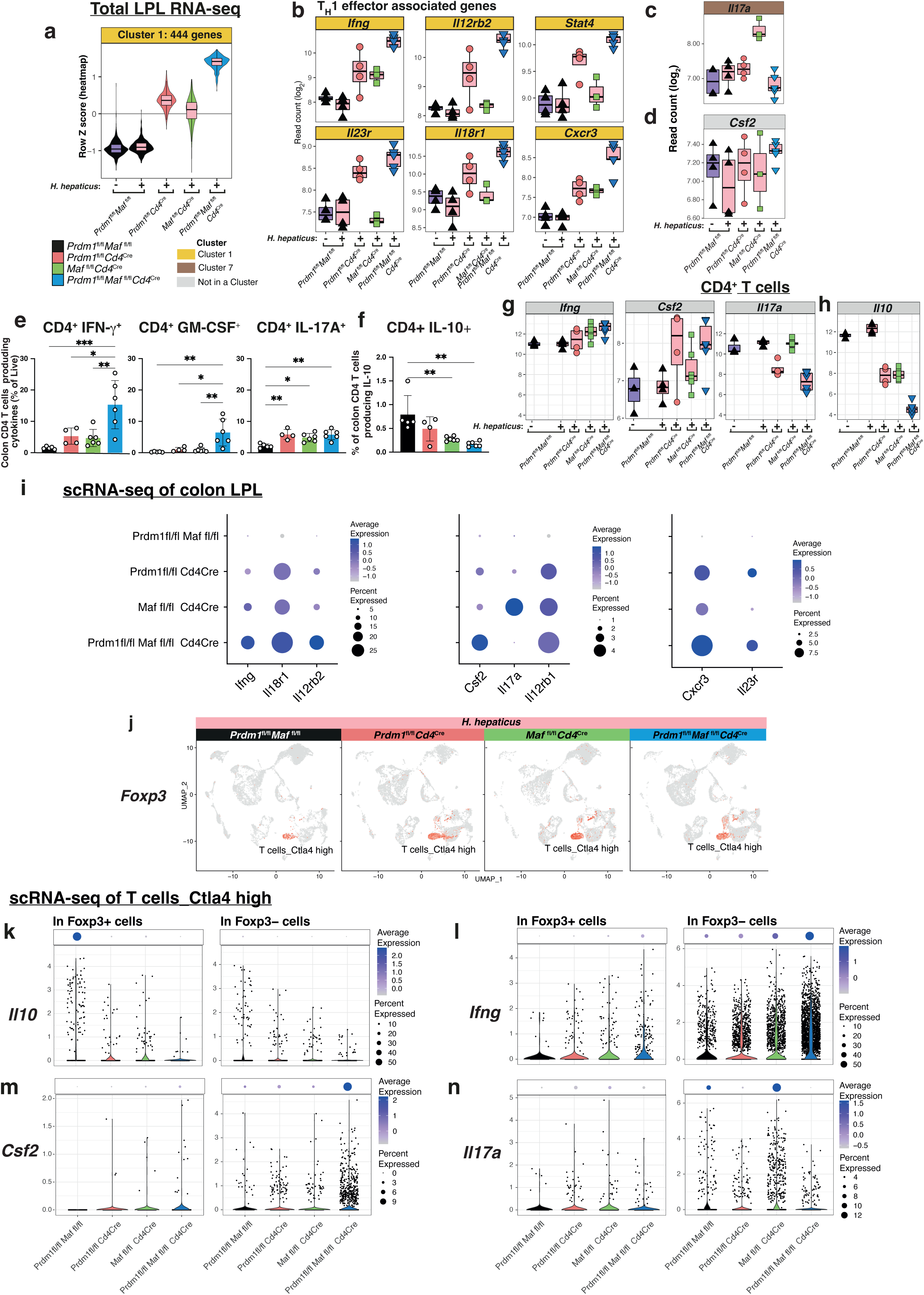
T cell-associated pro-inflammatory cytokines are increased in expression in *H. hepaticus* infected mice with *Cd4*^Cre^-mediated deletion of both *Prdm1* and *Maf* when compared to mice with deletion of either *Prdm1* or *Maf* and uninfected controls. **a)** Violin plots summarizing the expression values (quantified as z-scores) of the genes in Cluster 1 (**Fig.1f**) of the bulk tissue LPL RNA-seq. **b-d)** Boxplots of log2(normalized read counts) of **b)** representative genes in Cluster 1, c) *Il17a* (Cluster7) and d) *Csf2*. **e-f)** Flow cytometry analysis of colon lamina propria CD4+ T cells (Live CD90.2+ TCR-β+ CD4+ CD8-), **e)** as percentages from live cells, producing IFN-γ, GM-CSF, and IL-17A; and **f)** percentage of CD4+ T cells producing IL-10. Each dot within the barplots represents an individual mouse analyzed. Graph shows mean ±s.d., analyzed by one-way ANOVA followed by Tukey’s post-hoc test (*=p value ≤ 0.05, **=p value ≤ 0.01, ***=p value ≤ 0.001, ****=p value ≤ 0.0001). Data from n=3-6 mice. **g-h)** RNA-seq was performed on sorted CD4+ T cells from colon LPL isolated from uninfected *Prdm1*^fl/fl^*Maf*^fl/fl^ controls and *H. hepaticus* infected *Prdm1*^fl/fl^*Maf*^fl/fl^ mice and mice with *Cd4*^Cre^-mediated deletion of either *Prdm1*, *Maf*, or both *Prdm1* and *Maf*. Boxplots of log2(normalized read counts) of g) *Ifng, Csf2* and *Il17a* and h) *Il10*. **i)** Dotplot of scRNA-seq gene expression of selected genes *Ifng, Il18r1, ll12rb2, Csf2, Il17a, Il12rb1, Cxcr3* and *Il23r* in colon LPLs from *H. hepaticus* infected control *Prdm1*^fl/fl^*Maf*^fl/fl^ mice and mice with *Cd4*^Cre^-mediated deletion of either *Prdm1*, *Maf*, or both *Prdm1* and *Maf*. The dot size represents the percentage of cells per cell cluster expressing the gene in question and the expression level indicated by the colour-scale. **j)** Expression of *Foxp3*, as assessed by scRNA-seq within the UMAP visualization of annotated scRNA-seq datasets, within each of *H. hepaticus* infected conditions. **k-n)** Expression at the single cell level of k) *Il10*, l) *Ifng*, m) *Csf2* and n) *Il17a* in cluster Ctla4 high cells expressing *Foxp3* (Foxp3+) or not (Foxp3-). The distribution of expression in cells within each condition is shown by the violin plots and the dotplot (top panels) displays the proportion of cells expressing the gene in question.

### T cell-specific deletion of *Prdm1* and *Maf* disrupts different effector T cell signatures while both transcription factors co-operate to induce *Il10*

Quantitation of the total number of DEGs in the ‘T cell activation/T helper cells’, αβ-/γ8-T cells and ILC Cluster 1 from the bulk tissue RNA-Seq data (Fig.1f,g) confirmed that they were the most highly expressed in the LPLs from double *Prdm1*^fl/fl^*Maf*^fl/fl^*Cd4*^Cre^ mice infected with *H. hepaticus* and to a much lesser extent in each of the single *Prdm1^fl/fl^Cd4^Cre^*and *Maf^fl/fl^Cd4^Cre^* infected mice, as compared to control infected and uninfected mice which showed no quantitative differences in gene expression (Fig. 3a,b). The increased DEGs in Cluster 1, were dominated by increased levels of effector genes attributable to activated T cells, TH1 effector cells and NK/ILC1 cells, including *Ifng, Il12rb2, Il23r, Il18r1, Stat4, and Cxcr3* (Fig. 3b and Extended Supplementary Table 3). Strikingly, increased expression of *Il17a* was observed only in the LPL from *H. hepaticus* infected *Maf^fl/fl^Cd4^Cre^*mice as compared with controls (Fig. 3c). Expression of *Csf2* was increased in the LPL from the *H. hepaticus* double deficient *Prdm1*^fl/fl^*Maf*^fl/fl^*Cd4*^Cre^ mice (Fig. 3d). These findings were validated by ICS and flow cytometry (Fig. 3e; Extended Fig. 4a-h). Increased IFN-γ- producing CD4^+^ T cells and IFN-γ-producing total lymphocytes were observed in LPL from all the *H. hepaticus* infected T cell-specific transcription factor deficient mice, with maximal increases observed in the infected double deficient *Prdm1*^fl/fl^*Maf*^fl/fl^*Cd4*^Cre^ mice (Fig. 3e; Extended Fig.4c-f). A striking increase in GM-CSF protein production was observed in IFN-γ-producing CD4^+^ T cells from the LPL of infected double deficient *Prdm1*^fl/fl^*Maf*^fl/fl^*Cd4*^Cre^ mice only, accounting for the biggest increase in GM-CSF producing cells in the LPL (Fig. 3e; Extended Fig. 4e-h). Production of IL-17 protein was found to be increased in the LPL from the different T-cell-specific transcription factor deficient mice upon *H. hepaticus* infection (Fig. 3e; Extended Fig. 4e-h).

**Fig. 4.**
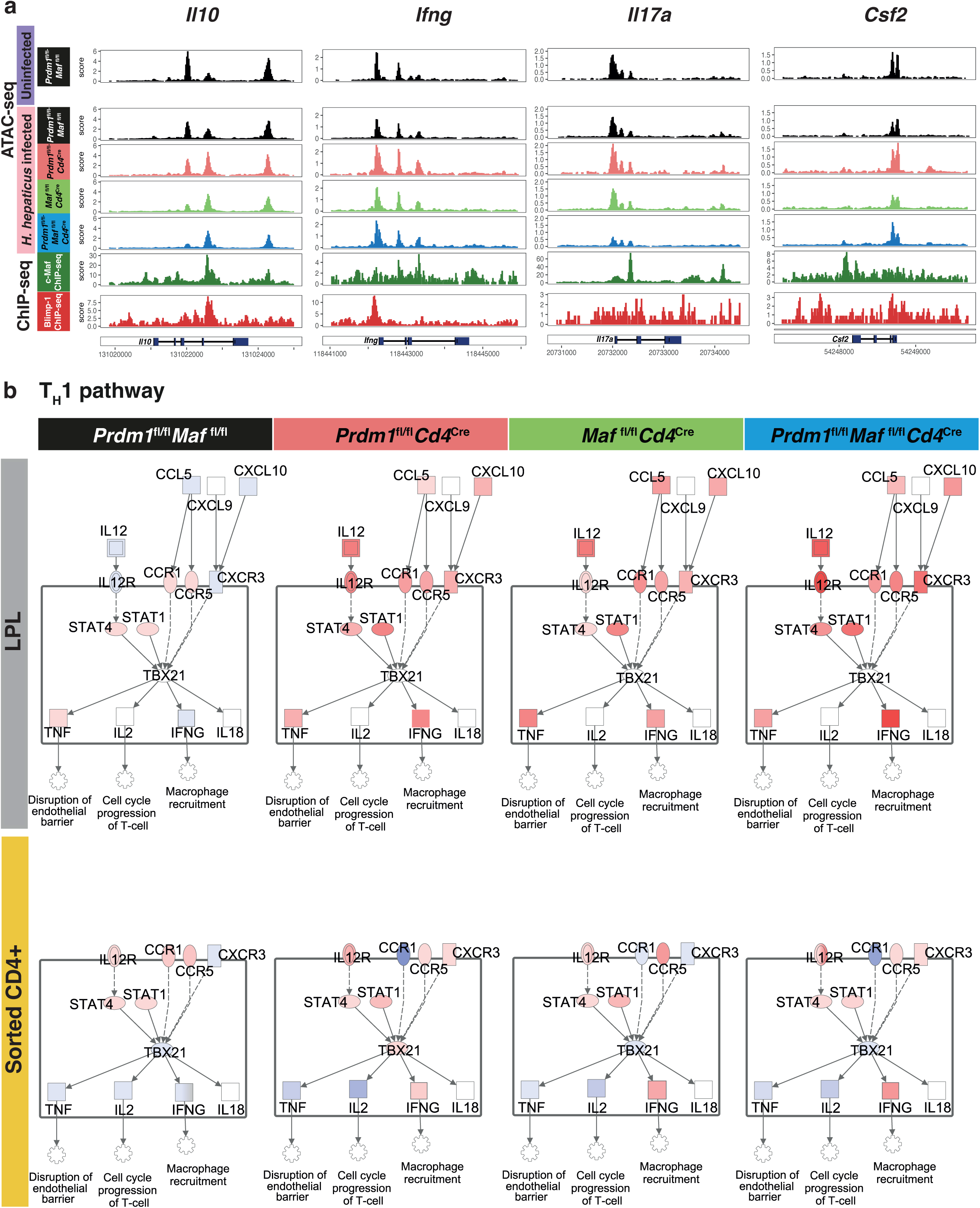
Integration of multi-omic datasets identifies the binding sites of Blimp-1 and/or c-Maf at *Il10, Ifng, Il17a* and *Csf2* loci and upregulation of the Th1 pathway in *H.* hepaticus infected mice with *Cd4*^Cre^-mediated deletion of both *Prdm1* and *Maf* when compared to mice with deletion of either *Prdm1* or *Maf* and uninfected controls. **a)** ATAC-seq was performed on sorted CD4+ T cells from colon LPL isolated from uninfected *Prdm1*^fl/fl^*Maf*^fl/fl^ control and *H. hepaticus* infected *Prdm1*^fl/fl^*Maf*^fl/fl^ mice and mice with *Cd4*^Cre^-mediated deletion of either *Prdm1*, *Maf*, or both *Prdm1* and *Maf*. Genome browser tracks of ATAC-seq data from each condition together with publicly available c-Maf (green) and Blimp-1 (red) ChIP-seq datasets in the *Il10*, *Ifng*, *Il17a* and *Csf2* loci. **b)** IPA was used to overlay bulk tissue LPL RNA-seq (top panel) and sorted CD4+ T cells RNA-seq (bottom panel) onto the Th1 pathway. Gene expression fold- changes in each *H. hepaticus* infected conditions relative to the uninfected *Prdm1*^fl/fl^*Maf*^fl/fl^ control were overlayed onto the “Th1 pathway”. A fixed scale of -5 (blue) to 3.5 (red) was kept between all conditions.

Analysis of RNA-Seq data from flow cytometry purified CD4^+^ T cells from LPLs from *H. hepaticus* infected *Prdm1^fl/fl^Cd4^Cre^*mice and *Maf*^fl/fl^*Cd4*^Cre^ mice, showed a similar increase in expression of *Ifng* and *Csf2* activated TH1 effector cells, with the greatest increase again observed in the double deficient *Prdm1^fl^*^/fl^*Maf*^fl/fl^*Cd4*^Cre^ (Fig. 3g; Extended Data Supplementary Table 4). However, the relative increase in *Ifng* expression as compared to controls was not as pronounced as in total LPL (Fig. 3b), suggesting that increased numbers of CD4^+^ T cells in the LPL of the *H. hepaticus* infected T cell-specific transcription factor deficient mice are contributing to the global increased levels of *Ifng* expression Fig. 3b). Additionally, other sources of *Ifng* such as NK/ILC1 cells could potentially play a major role in this response, as shown by the scRNA-Seq data (Fig. 2e; Extended Data Supplementary Table 4). The level of *Il17a* expression in purified CD4^+^ T cells from the LPL was again highest in the infected *Maf*^fl/fl^*Cd4*^Cre^ mice but in this case was at a similar level to that in uninfected and infected control mice suggesting that microbiota may be maintaining *Il17a* expression in T cells (Fig. 3g), as previously suggested. However, the expression of *Il17a* was much decreased in purified CD4^+^ T cells from LPL from infected double deficient *Prdm1^fl^*^/fl^*Maf*^fl/fl^*Cd4*^Cre^ and *Prdm1^fl/fl^Cd4^Cre^* mice, suggesting that genes negatively regulated by Blimp1 may actively repress the expression of *Il17a* in CD4^+^ T cells (Fig. 3g).

scRNA-seq analysis showed that levels of expression and percentage of cells expressing *Ifng, Il18r1, Il12rb2, Csf2, Il12rb1, Cxcr3* and *Il23r* were highest in the LPL from *H. hepaticus* infected double deficient *Prdm1^fl^*^/fl^*Maf*^fl/fl^*Cd4*^Cre^, and to a much lesser extent in both the *Prdm1^fl/fl^Cd4^Cre^* mice and *Maf*^fl/fl^*Cd4*^Cre^ mice, (Fig. 3i). This suggests that *Prdm1* and *Maf* not only regulate gene expression of these Type 1 associated molecules but also the abundance of these effector cells and that both transcription factors cooperate to control Type 1 responses. Conversely, scRNA-seq data showed exclusive detection of *Il17a* at the level of expression and percentage of *Il17a* expressing cells in LPLs from *H. hepaticus* infected *Maf*^fl/fl^*Cd4*^Cre^ mice, suggesting that c-Maf not only controls expression of *Il17a* but additionally the numbers of *Il17a* expressing cells (Fig. 3i).

IL-10 protein production was reduced in each of the *Prdm1^fl/fl^Cd4^Cre^*and *Maf^fl/fl^Cd4^Cre^* LPL and completely diminished in the CD4^+^ T cells from the infected double deficient *Prdm1*^fl/fl^*Maf*^fl/fl^*Cd4*^Cre^ mice as compared to infected control mice (Fig. 3f). In keeping with the ICS IL-10 protein data, the expression of *Il10* was diminished in purified CD4^+^ T cells from the LPL of infected *Prdm1^fl/fl^Cd4^Cre^* and *Maf*^fl/fl^*Cd4*^Cre^ mice, and to the greatest extent in the double deficient *Prdm1^fl^*^/fl^*Maf*^fl/fl^*Cd4*^Cre^ mice (Fig. 3h; Extended Supplementary Table 5).

ScRNA-seq data was then further interrogated to identify the source of cells expressing *Il10* and proinflammatory cytokines in the LPLs from the *H. hepaticus* infected T cell-specific transcription factor deficient mice. Foxp3^+^ regulatory T cells (Tregs), contained within the Ctla4 high cluster of T cells (Fig. 3j), were identified as the main *Il10* expressing T cells (Fig. 3k). Expression *Il10* and percentage of *Il10* producing cells by scRNA-Seq was found to be diminished in both Foxp3^+^ Tregs from the LPL of *H. hepaticus* infected *Prdm1^fl/fl^Cd4^Cre^*, *Maf*^fl/fl^*Cd4*^Cre^ and to the greatest extent in the double deficient *Prdm1^fl^*^/fl^*Maf*^fl/^fl*Cd4*^Cre^, as compared to infected control mice, and similarly in the very low numbers of Foxp3^-^ CD4^+^ T cells (Fig. 3k). Flow cytometric analysis revealed however, that Foxp3^+^ Tregs showed a graded increase in numbers in the LPL from *H. hepaticus* infected *Prdm1^fl/fl^Cd4^Cre^*, *Maf*^fl/fl^*Cd4*^Cre^ mice and double deficient *Prdm1^fl^*^/fl^*Maf*^fl/fl^*Cd4*^Cre^ mice respectively (Extended Fig.4j,k). In contrast, Foxp3^+^RORγt^+^ T cells were almost completely abolished in the LPL from infected *Maf*^fl/fl^*Cd4*^Cre^ mice (Extended Fig. 4j) in keeping with a previous report^31^, whilst increased in LPL from infected *Prdm1^fl/fl^Cd4^Cre^* and to a lesser extent double deficient *Prdm1^fl^*^/fl^*Maf*^fl/fl^*Cd4*^Cre^ mice as compared to control mice (Extended Fig. 4j,k). Despite this, the LPL from the infected double deficient *Prdm1^fl^*^/fl^*Maf*^fl/^fl*Cd4*^Cre^ mice expressed the lowest levels of *Il10* (Fig. 3f,h,k) and exhibited the maximum pathology (Fig. 1b,c; Fig. 2b).

In contrast to the dominance of *Il10* expression in Foxp3^+^ Tregs in the Ctla4 high cluster of T cells, the expression of *Ifng* and *Csf2* was highest in the Foxp3^-^ Ctla4 high T cell cluster in the LPLs from *H. hepaticus* infected mice and scarcely detectable in Foxp3^+^ Tregs (Fig. 3l,m). Both cytokines showed a graded increase in the LPLs from infected *Prdm1^fl/fl^Cd4^Cre^*and *Maf*^fl/fl^*Cd4*^Cre^ mice, with the highest level in double deficient *Prdm1^fl^*^/fl^*Maf*^fl/fl^*Cd4*^Cre^ mice respectively (Fig. 3kl,m). Conversely, *Il17a* expression by scRNA-Seq, was largely in Foxp3^-^ CD4^+^ T cells and was highest in LPL from infected *Maf*^fl/fl^*Cd4*^Cre^ mice (Fig. 3n), in keeping with data from the RNA-Seq from flow cytometry purified CD4+ T cells (Fig. 3g). This diminished expression of *Il17a* in LPLs from infected *Prdm1^fl/fl^Cd4^Cre^*and *Prdm1^fl^*^/fl^*Maf*^fl/^fl*Cd4*^Cre^ mice again suggest repression of the *Il17a* response by Blimp-1 regulated factors (Fig. 3n). Collectively these data suggest that the intestinal pathology resulting from T cell-specific deficiency in *Maf* differs qualitatively from that of *Prdm1* and even *Prdm1/Maf* double deficiency due to an IL-17-mediated effector cytokine response to *H. hepaticus* infection, as well an increased Type 1 mediated response controlled by both transcription factors.

A combination of ATAC-Seq and RNA-Seq data from purified CD4^+^ T cells from colon LPL from the *H. hepaticus* infected T cell-specific transcription factor deficient mice was integrated with published ChIP-Seq data and revealed direct common and distinct binding sites for Blimp-1 and c-Maf in the *Ifng, Il17a* and *Csf2* loci and the *Il10* locus (Fig. 4a), in addition to those previously reported for *Il10*^14^. Our data are supportive of cooperative and independent roles of these transcription factors in the positive regulation of *Il10* and additionally in negative regulation of proinflammatory cytokine genes to ensure strict control of the immune response to pathogens and pathobionts. Pathway analysis applied to the bulk tissue RNA-Seq data revealed an increase in IL-12 and downstream Stat1/Stat4 signalling for *Ifng* induction in the LPL and CD4^+^ T cells from the *H. hepaticus* infected *Prdm1^fl/fl^Cd4^Cre^*, *Maf*^fl/fl^*Cd4*^Cre^ and double deficient *Prdm1^fl^*^/fl^*Maf*^fl/fl^*Cd4*^Cre^ mice respectively, with the biggest increase observed in the double deficient mice, as compared to infected control mice (Fig. 4b). This again supports direct action of Blimp-1 and c-Maf on *Ifng* expression (Fig. 4b). *IL- 12b/IL-23a* and *IL-21* expression was increased in the LPL from all *H. hepaticus* infected *Prdm1^fl/fl^Cd4^Cre^*, *Maf*^fl/fl^*Cd4*^Cre^ and double deficient *Prdm1^fl^*^/fl^*Maf*^fl/fl^*Cd4*^Cre^ mice as compared to infected control mice (Extended Fig. 5). While pathway analysis showed that downstream expression of IL-22 was also similarly increased in the LPL from all the infected T cell specific transcription factor-deficient mice, downstream *IL- 17a* expression was increased to the highest level in the LPL from *Maf*^fl/fl^*Cd4*^Cre^ mice, and actually decreased in the LPL from the double deficient *Prdm1^fl^*^/fl^*Maf*^fl/fl^*Cd4*^Cre^ mice (Extended Fig. 5). This decrease in *IL-17a* expression observed by pathway analysis was more marked in purified CD4^+^ T cells from the LPL of infected *Prdm1^fl/fl^Cd4^Cre^* and double deficient *Prdm1^fl^*^/fl^*Maf*^fl/fl^*Cd4*^Cre^ mice as compared to the infected *Maf*^fl/fl^*Cd4*^Cre^ and control infected mice, supporting active repression of the *Il17a* gene (Extended Fig. 5). Expression of *Csf2* observed by pathway analysis was again increased in CD4^+^ T cells from LPL of all the *H. hepaticus* infected *Prdm1^fl/fl^Cd4^Cre^*, *Maf*^fl/fl^*Cd4*^Cre^ and *Prdm1^fl^*^/fl^*Maf*^fl/fl^*Cd4*^Cre^ mice as compared to control infected mice (Extended Fig. 5).

**Fig. 5.**
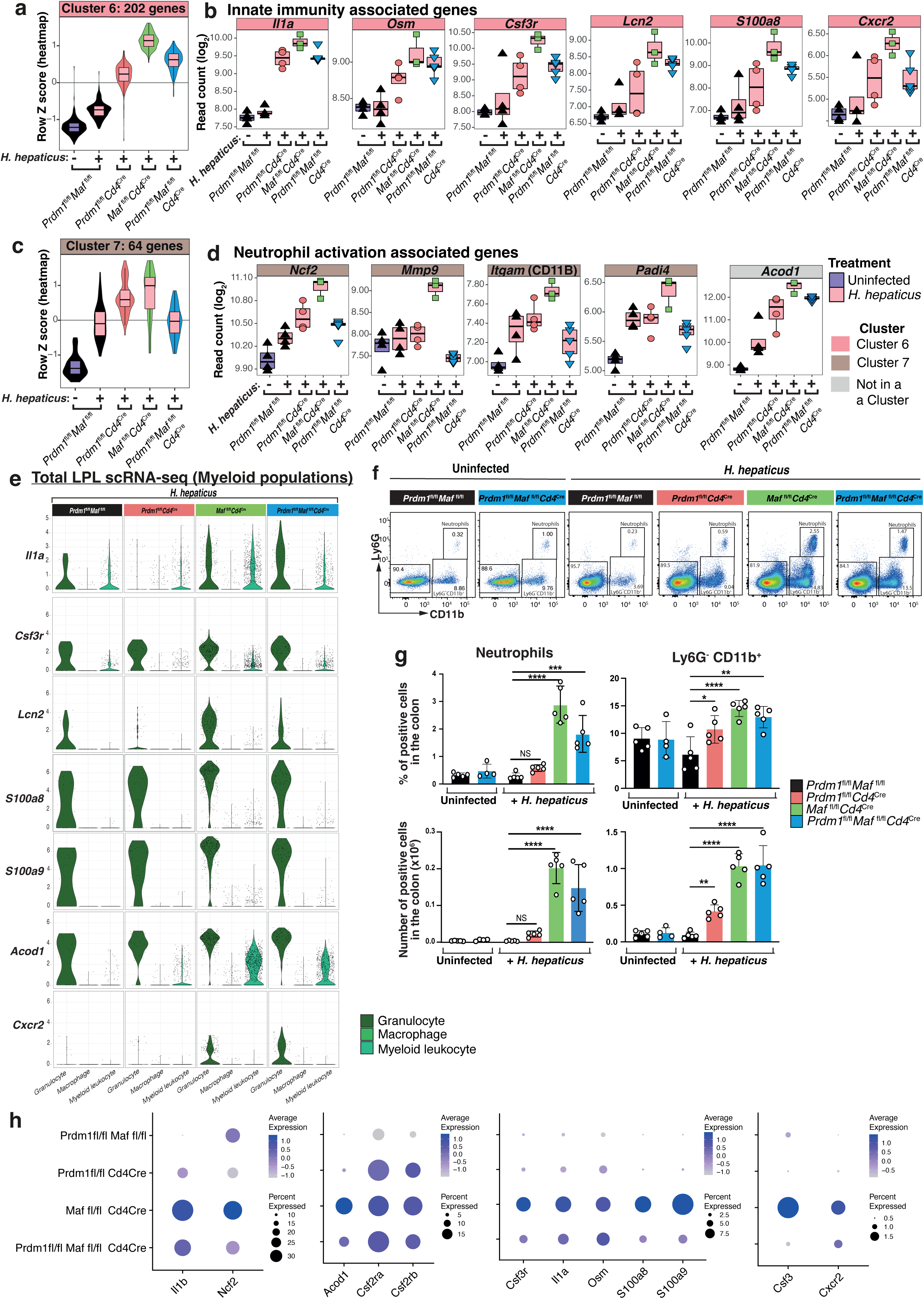
Granulocyte-associated genes are increased in expression in *H. hepaticus* infected mice with *Cd4*^Cre^-mediated deletion of *Maf* when compared to infected mice with deletion of either *Prdm1* or both *Prdm1* and *Maf* and uninfected controls. a-d) Violin plots summarizing the expression values (quantified as z-scores) and boxplots of example genes of **a-b)** Innate immunity associated genes found in Cluster 6 and **c-d)** Neutrophil activation associated genes found in Cluster 7 of the bulk tissue LPL RNA-seq data analysis from **Fig.1f**. **e)** Expression of selected innate immunity and granulocyte-associated genes queried in the Granulocyte, Macrophage and Myeloid leukocyte cell clusters across all infected *H. hepaticus* infected conditions within the colon LPL scRNA-seq dataset. **f)** Representative flow plots and gating strategy used for the flow cytometry analysis of neutrophils (Live CD90.2- TCR-β- CD19-CD11b+ Ly6G+) across uninfected *Prdm1*^fl/fl^*Maf*^fl/fl^ and *Prdm1*^fl/fl^*Maf*^fl/fl^*Cd4*^Cre^, and *H. hepaticus* infected *Prdm1*^fl/fl^*Maf*^fl/fl^ mice and mice with *Cd4*^Cre^-mediated deletion of either *Prdm1*, *Maf*, or both *Prdm1* and *Maf*. **g)** Barplots of percentage from live (top panels) and absolute cell counts (bottom panels) of neutrophils and Ly6G-CD11b+ cells from lamina propria of the colon. Each dot within the barplots represents an individual mouse analyzed. Graph shows mean ±s.d., analyzed by one-way ANOVA followed by Tukey’s post-hoc test (*=p value ≤ 0.05, **=p value ≤ 0.01, ***=p value ≤ 0.001, ****=p value ≤ 0.0001). Data from n=4-5 mice. **h)** Dotplot of scRNA-seq gene expression of selected genes *Il1b, Ncf2, Acod1, Csf2ra, Csf2rb, Csf3r, Il1a, Osm, S100a8, S100a9, Csf3 and Cxcr2,* in the colon LPLs from *H. hepaticus* infected control *Prdm1*^fl/fl^*Maf*^fl/fl^ mice and mice with *Cd4*^Cre^-mediated deletion of either *Prdm1*, *Maf*, or both *Prdm1* and *Maf*. The dot size represents the percentage of cells per cell cluster expressing the gene in question and the expression level indicated by the colour-scale.

### T cell-derived Blimp-1 and c-Maf control myeloid cells and innate immunity

The average increased expression of ‘innate immunity and myeloid-associated’ genes in the bulk tissue RNA-Seq data, Cluster 6 (Fig. 1f,g) from the LPL from *H. hepaticus* infected *Maf*^fl/fl^*Cd4*^Cre^ and to a lesser extent *Prdm1*^fl/fl^*Cd4*^Cre^ and double deficient *Prdm1*^fl/fl^*Maf*^fl/fl^*Cd4*^Cre^ mice as compared to controls, was confirmed quantitatively (Fig. 5a), for example for *Il1a* and *Osm,* encoding Oncostatin M (Fig. 5b), previously associated with colitis^11, 32, 33^. Granulocyte-associated genes in this cluster including, *Csf3r, Lcn2, S100a8* and *Cxcr2,* were most highly expressed in the LPL from *Maf*^fl/fl^*Cd4*^Cre^ mice and to a much lesser extent in the double deficient *Prdm1*^fl/fl^*Maf*^fl/fl^*Cd4*^Cre^ and *Prdm1*^fl/fl^*Cd4*^Cre^ infected mice, as compared to controls (Fig. 5b). Similarly, while the average gene expression in the bulk tissue RNA-Seq data granulocyte Cluster 7 (Fig. 1f,g) was increased in the LPL of control mice upon infection with *H. hepaticus* a further increase in the *Maf*^fl/fl^*Cd4*^Cre^ and to a lesser extent in the *Prdm1*^fl/fl^*Cd4*^Cre^ infected mice, but not in the double deficient *Prdm1*^fl/fl^*Maf*^fl/fl^*Cd4*^Cre^ infected mice, which showed similar levels to those of the *H. hepaticus* infected control mice (Fig. 5c). Genes associated with granulocyte/neutrophil activation including, *Ncf2*, *Itgam, Mmp9* and *Padi4,* showed the highest expression in the LPL from infected *Maf*^fl/fl^*Cd4*^Cre^ mice but not in the *Prdm1*^fl/fl^*Cd4*^Cre^ or double deficient *Prdm1*^fl/fl^*Maf*^fl/fl^*Cd4*^Cre^ infected mice, which showed similar or reduced levels to those from infected control mice, the latter suggesting potential counter-regulatory mechanisms provided by Blimp-1 signalling in T cells (Fig. 5d). Additionally, *Acod1*, a gene encoding enzyme aconitate decarboxylase 1 (Irg1) which produces the metabolite itaconate in myeloid cells^34, 35, 36^ was found to be most highly expressed in LPL from infected *Maf*^fl/fl^*Cd4*^Cre^ mice and double deficient *Prdm1*^fl/fl^*Maf*^fl/fl^*Cd4*^Cre^ infected mice (Fig. 5d). scRNA-Seq analysis showed that the granulocyte/neutrophil and myeloid leukocyte clusters were the source of *Il1a* and *Acod1* in the LPL from infected *Maf*^fl/fl^*Cd4*^Cre^ and double deficient *Prdm1*^fl/fl^*Maf*^fl/fl^*Cd4*^Cre^ mice (Fig. 5e). The expression of genes associated with granulocytes/neutrophils, including *Lcn2, S100a8, S100a9* and *Cxcr2* was exclusively detected in the ‘granulocyte/neutrophil’ cluster by scRNA-Seq (Fig. 5e). Although basal expression of some of these genes was observed in the low number of granulocytes detected by scRNA-Seq in the infected control mice, increased expression of these neutrophil associated genes (Fig. 5e) and granulocyte percentage and numbers (Extended Fig. 2b,c) were observed in the LPL from the *Maf^fl/fl^Cd4^Cre^*and the double deficient *Prdm1^fl/fl^Maf^fl/fl^Cd4^Cre^*infected mice. In keeping with these findings by scRNA-Seq, increased percentage and numbers of Ly6G^+^CD11b^+^ neutrophils cells were observed by flow cytometry analysis, in the LPL from the *Maf*^fl/fl^*Cd4*^Cre^ but to a lesser extent in double *Prdm1*^fl/fl^*Maf*^fl/fl^*Cd4*^Cre^ *H. hepaticus* infected mice, as compared to uninfected or infected control mice and infected *Prdm1*^fl/fl^*Cd4*^Cre^ mice where these populations were barely detectable (Fig. 5f,g). This was reinforced by analysis of combined gene expression and percentage in the scRNA-seq data for granulocyte-specific genes, such as *Ncf2, Csf3r, S100a8* and *S100a9,* as well as innate cells genes *Il1a, Il1b* and *Acod1,* which were maximal in the LPLs from the infected *Maf*^fl/fl^*Cd4*^Cre^ mice (Fig. 5h). On the other hand, Ly6G^-^CD11b^+^ myeloid cells, while present in uninfected and infected control mice at low numbers, showed a significant increase by flow cytometry in the LPL from the *H. hepaticus* infected *Prdm1*^fl/f^*Cd4*^Cre^ mice, and to the greatest extent in both *Maf*^fl/fl^*Cd4*^Cre^ and double deficient *Prdm1^fl^*^/fl^*Maf*^fl/fl^*Cd4*^Cre^ mice as compared to the uninfected or infected control mice (Fig. 5 f,g). This increase in myeloid cells is in keeping with increased expression of *Il1a, Il1b, Acod1* and the *Csf2ra* and *Csf2rb* in the LPLs from both the infected *Maf*^fl/fl^*Cd4*^Cre^ and double deficient *Prdm1^fl^*^/fl^*Maf*^fl/fl^*Cd4*^Cre^ mice (Fig. 5e,h).

### Increased T cell-myeloid cell interactions in LPL from *H. hepaticus* infected T-cell-specific *Prdm1* and *Maf* deficient mice

Single cell gene expression analysis of ligand-receptor pairs using CellChat^37^ was used to infer putative cell-to-cell crosstalk (Fig. 6; Extended Fig. 6). Strong stromal/epithelial to B cell interactions were apparent in the LPL from *H. hepaticus* infected control mice which were significantly decreased in those from *Prdm1*^fl/f^*Cd4*^Cre^, *Maf*^fl/fl^*Cd4*^Cre^ and double deficient *Prdm1^fl/fl^Maf^fl/fl^Cd4^Cre^*mice which instead showed strong outgoing and incoming interactions between the T cell_Ctla4 high cluster cell and innate cell interactions (Extended Fig. 6a-c). Analysis of the immune cells of interest from this study, revealed outgoing and incoming interactions between the T cell_Ctla4 high population and B cells in the LPL from *H. hepaticus* infected control mice, while in the *Prdm1*^fl/f^*Cd4*^Cre^, *Maf*^fl/fl^*Cd4*^Cre^ and double deficient *Prdm1^fl/fl^Maf^fl/fl^Cd4^Cre^*mice myeloid leukocyte/macrophage populations delivered increased strength and number of signals to the T cell_Ctla4 high population, which in turn signalled back to these myeloid cells (Fig. 6a-c). These findings were validated by immunofluorescent staining of colon sections (Fig. 6d; Extended Fig. 7). CD4^+^ T cells (white) and CD68^+^ mononuclear phagocytes (magenta) were found to be increased and in close proximity in *H. hepaticus* infected *Prdm1*^fl/f^*Cd4*^Cre^ mice, and to a greater extent in *Maf*^fl/fl^*Cd4*^Cre^ and double deficient *Prdm1*^fl/fl^*Maf*^fl/fl^*Cd4*^Cre^ infected mice, as compared to control mice (Fig. 6d). Neutrophils/granulocytes staining positive for MPO (green) were found in colon sections from *Maf*^fl/fl^*Cd4*^Cre^ and double deficient *Prdm1*^fl/fl^*Maf*^fl/fl^*Cd4*^Cre^ infected mice (Fig. 6d; Extended Fig. 7), in keeping with the bulk tissue RNA-Seq, scRNA-Seq and flow cytometry data described above (Fig. 1,2 and 5) and appeared to co-localise for the most part with CD68^+^ mononuclear phagocytes and CD4^+^ T cells (Fig. 6d). However, the interactions between neutrophils and these cell types were relatively weakly detected by CellChat as compared with those between myeloid cells and T cells (Fig. 6b).

**Fig. 6.**
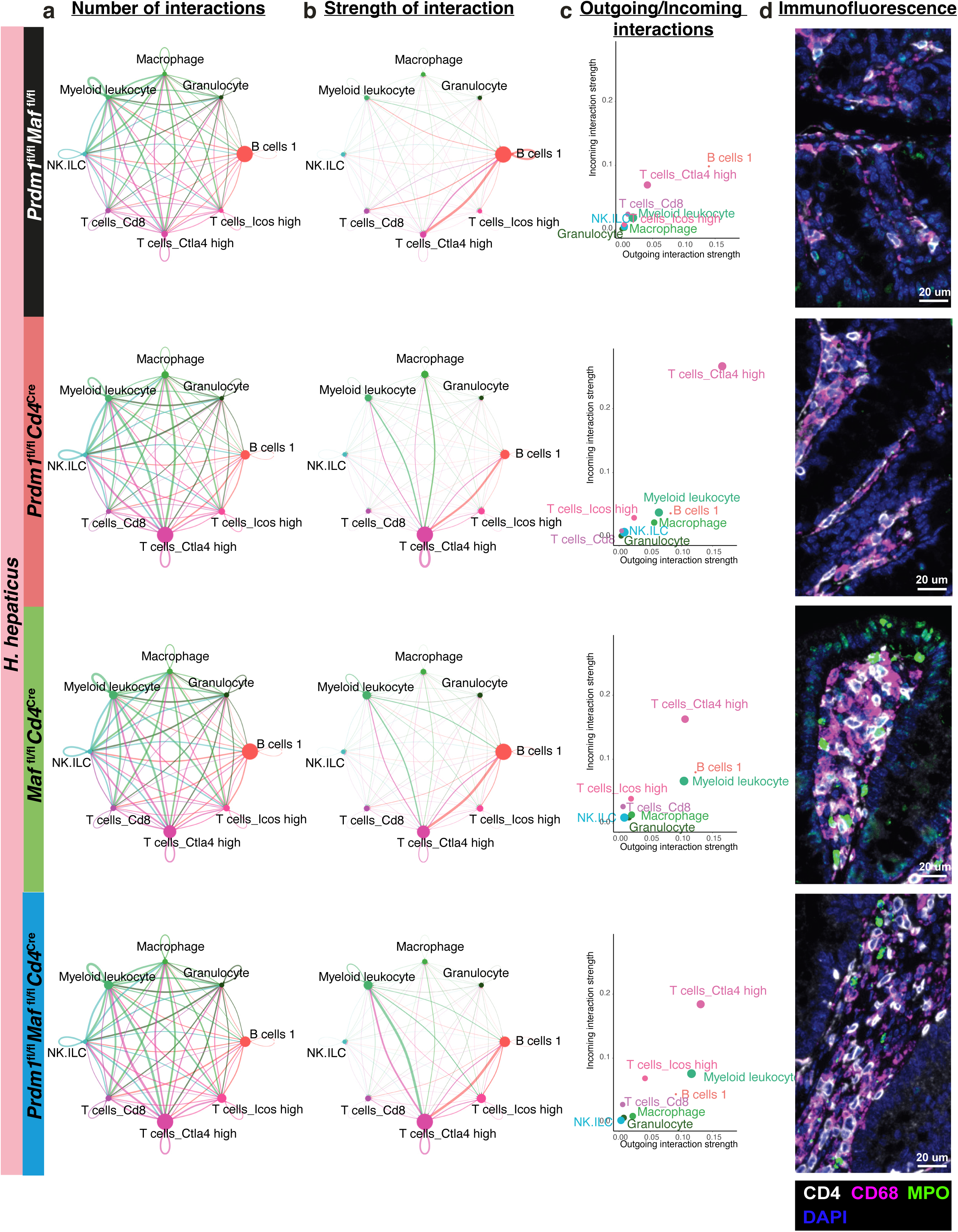
Inference of cell-to-cell communication networks reveals an increase in the interactions between T cells, macrophages and neutrophils in *H. hepaticus* infected mice with *Cd4*^Cre^-mediated deletion of *Prdm1*, *Maf,* or both *Prdm1* and *Maf* when compared to infected *Prdm1*^fl/fl^*Maf*^fl/fl^ mice. Cell-to-cell communication networks inferred using CellChat software from gene expression of ligands and their receptors in immune cell clusters of interest from the colonic LPL scRNA-seq dataset. **a)** Number and **b)** strength of interaction of cell-to-cell interactions, represented in the edge width, in and *H. hepaticus* infected *Prdm1*^fl/fl^*Maf*^fl/fl^ mice and mice with *Cd4*^Cre^- mediated deletion of either *Prdm1*, *Maf*, or both *Prdm1* and *Maf*. **c)** Plot of “Outgoing interaction strength“ against “Incoming interaction strength” across all *H. hepaticus* infected conditions. In **a-c)** node size is proportional to the number of cells in each experimental group, and the edges are colored based on the cell clusters expressing the outgoing signals. **d)** Representative images (n=4-5) of colon sections by immunofluorescence, staining for CD4+ T cells (CD4, white), mononuclear phagocytes (CD68, magenta), neutrophils (MPO, green) and nuclear staining (DAPI, blue) from *H. hepaticus* infected *Prdm1*^fl/fl^*Maf*^fl/fl^ mice and mice with *Cd4*^Cre^-mediated deletion of either *Prdm1*, *Maf*, or both *Prdm1* and *Maf*. Scale bar = 20μm.

**Fig. 7.**
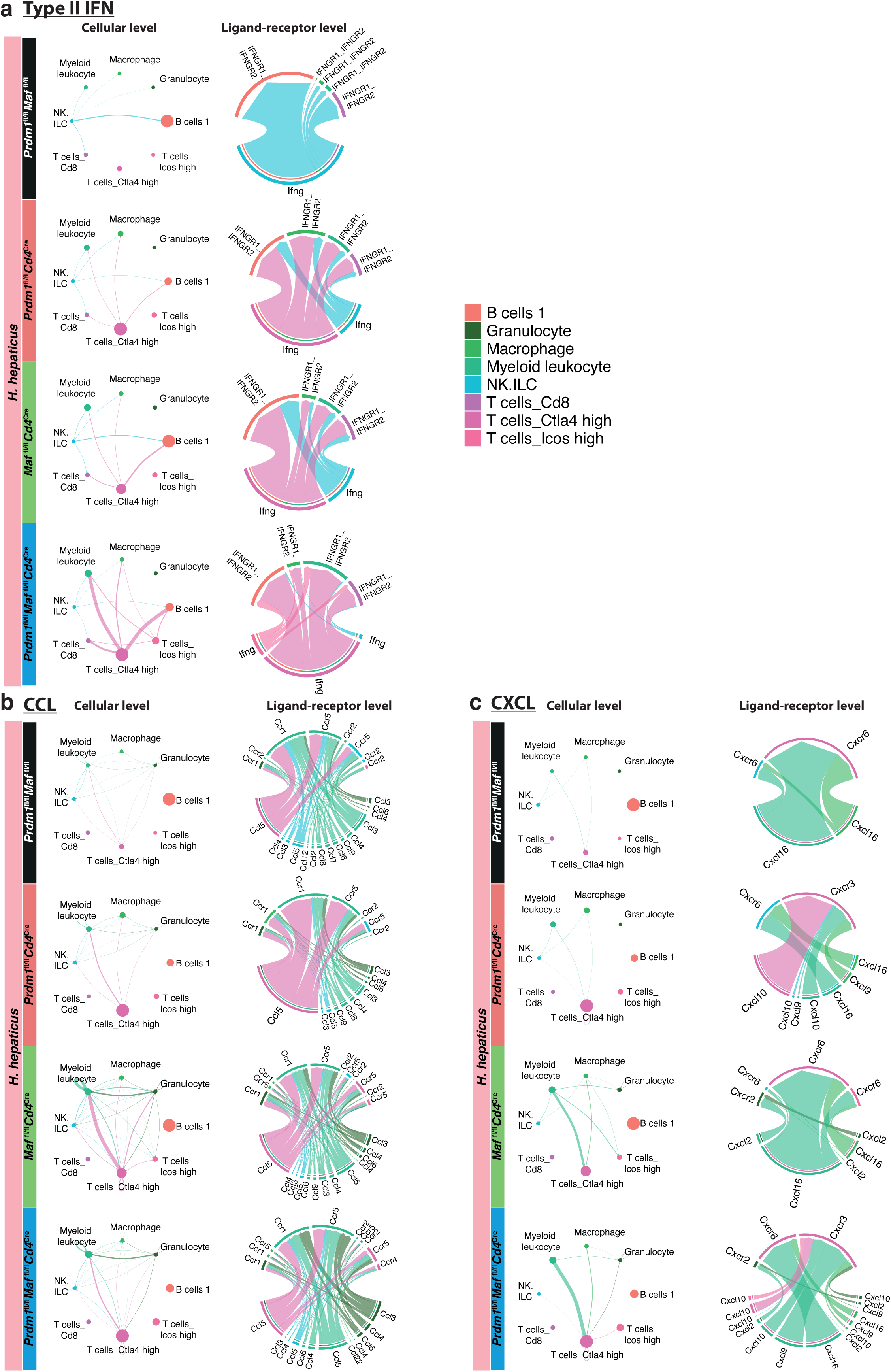
Putative cell-to-cell communication network analysis shows an increase in T cell-driven IFN-γ pathway and an increase in T cell, myeloid cell and granulocyte chemoattractant pathways in *H. hepaticus* infected mice with *Cd4*^Cre^-mediated deletion of *Prdm1*, *Maf,* or both *Prdm1* and *Maf* when compared to infected *Prdm1*^fl/fl^*Maf*^fl/fl^ control mice. Cell-to-cell communication networks underlying **a)** Type II IFN (IFN-γ), **b)** CCL and **c)** CXCL pathways across all *H. hepaticus* infected *Prdm1*^fl/fl^*Maf*^fl/fl^ mice and mice with *Cd4*^Cre^-mediated deletion of either *Prdm1*, *Maf*, or both *Prdm1* and *Maf*. The chord plot has receiver cells at the top (incoming signaling) and transmitter cells (outgoing signaling) the bottom. The edges are colored based on the cell clusters expressing the outgoing signals.

Having identified the major interacting cell types in the LPL we sought to understand the signalling pathways associated with the different cell types and how these signalling networks were impacted by T cell-specific deletion of *Prdm1, Maf* or both transcription factors in *H. hepaticus* infected mice (Extended Fig. 8). Increased outgoing and incoming signalling in myeloid leukocytes and T cell Ctla4 high cells was observed in *Prdm1*^fl/f^*Cd4*^Cre^, *Maf*^fl/fl^*Cd4*^Cre^ and double deficient *Prdm1*^fl/f^*Maf*^fl/f^*Cd4*^Cre^ mice (Extended Fig. 8). Strikingly, while IFN-II (IFN-γ) signalling in infected control mice was inferred to be delivered from NK cells to B cells, this was replaced by an outgoing signal from T cell Ctla4 high cells to incoming signalling to macrophages and myeloid leukocytes, as well as B cells, in the infected *Prdm1*^fl/f^*Cd4*^Cre^, *Maf*^fl/fl^*Cd4*^Cre^ and double deficient *Prdm1*^fl/f^*Maf*^fl/f^*Cd4*^Cre^ mice (Extended Fig. 8; Fig. 7a). Inferred outgoing and incoming *Csf1* signalling was exclusively detected between the myeloid populations in both *Maf*^fl/fl^*Cd4*^Cre^ and double deficient *Prdm1*^fl/f^*Maf*^fl/f^*Cd4*^Cre^ and weakly in the *Prdm1*^fl/f^*Cd4*^Cre^ infected mice (Extended Fig. 8 and 9), potentially contributing to the increased myeloid cells and neutrophils (Fig. 5g; Fig. 6d). The largest inferred increase in both outgoing and incoming signals observed in myeloid leukocytes was in the LPL from infected *Maf*^fl/fl^*Cd4*^Cre^ and double deficient *Prdm1*^fl/f^*Maf*^fl/f^*Cd4*^Cre^ mice (Extended Fig. 8), such as strong outgoing CCL and CXCL signalling (Extended Fig. 8). While the T cell Ctla4 high population were inferred to deliver a strong *Ccl5* signal to *Ccr1/Ccr5* on myeloid leukocytes (Fig. 7b), the myeloid leukocyte population delivered a strong *Cxcl16/Cxcl10* signal largely to *Cxcr6/Cxcr3* on the T cells_Ctla4 high population highlighting potential axes contributing to CD4 T cell accumulation in the LPLs of these mice (Fig. 7c). Consistent with the well-established role of *Cxcl2- Cxcr2* in neutrophil response, inferred interactions between the myeloid leucocyte population and granulocytes through the *Cxcl2-Cxcr2* axis, and additionally strong *Ccl* interactions within the myeloid leukocyte population and from myeloid leukocytes to granulocytes were detected, again only in the infected *Maf*^fl/fl^*Cd4*^Cre^ and double deficient *Prdm1*^fl/f^*Maf*^fl/f^*Cd4*^Cre^ mice (Fig. 7b,c), potentially contributing to the increased neutrophil and myeloid numbers discussed earlier (Fig. 5g; Fig. 6d). These analyses highlight dysregulated cytokine and chemokine cell-cell interaction networks downstream of disrupted transcriptional regulation in T cells by Blimp-1 and c-Maf that likely contribute to pathology during infection with a pathobiont.

**Fig. 8.**
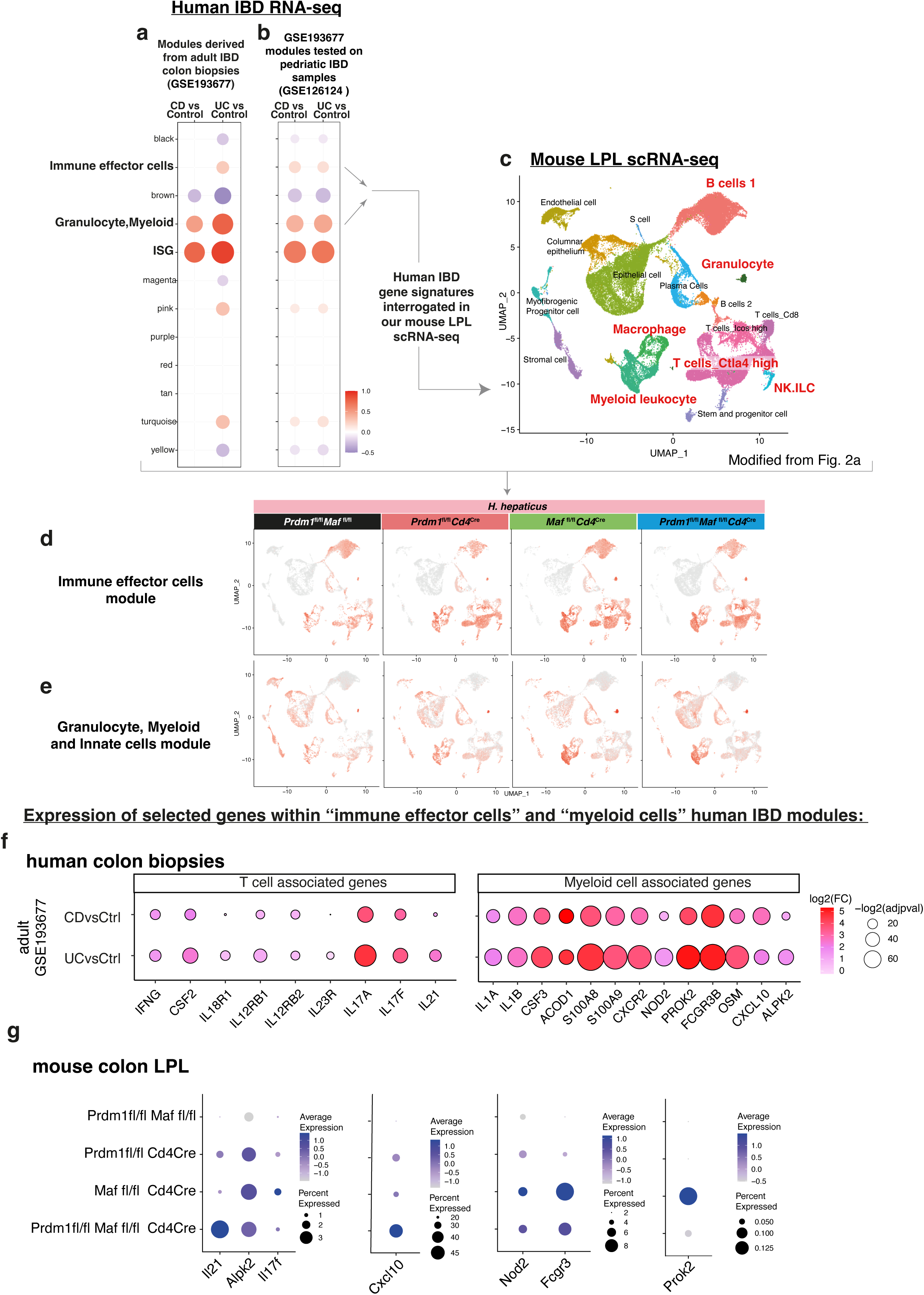
Analysis of human colon biopsies from IBD patients highlights shared immune cell and inflammation gene signatures between IBD patients and mice with *Cd4*^Cre^-mediated deletion of *Prdm1*, *Maf,* or both *Prdm1* and *Maf.* **a)** Modules of co-expressed genes were derived using WGCNA from human adult IBD colonic biopsies (GSE193677) and **b)** tested in an independent human pediatric IBD dataset (GSE126124). In the dotplots the color and size dots represent the fold enrichment in disease compared to control. Re-named modules indicate biological processes associated with the genes within a module. Fold-enrichment scores were derived using QuSAGE software, red and blue color indicated over- and under-abundance, respectively, of genes within a module (as compared to control samples). Size of the dots represents the relative degree of perturbation (larger dots represent a higher degree of perturbation), only modules with an adjusted p value<0.05 were considered significant and depicted in the plot. **a-c)** Enrichment of genes within the “Immune effector cells” and the “Granulocyte, Myeloid and Innate cells” modules were then tested in our mouse colon LPL scRNA-seq dataset. Scoring of the **d)** “Immune effector cells” and the **e)** “Granulocyte, Myeloid and Innate cells modules were projected into our scRNA-seq UMAP. **f)** Dotplot of over-expressed T cell or myeloid cell associated genes from human IBD colon biopsies found within the modules from Fig. 8a and b, against controls. The dot size represents the p value and the log2 fold change is indicated by the colour-scale. **g)** Dotplot of gene expression of selected genes found to be over-expressed in human IBD as in Fig. 8f, in colon LPLs scRNA-seq from *H. hepaticus* infected control *Prdm1*^fl/fl^*Maf*^fl/fl^ mice and mice with *Cd4*^Cre^-mediated deletion of either *Prdm1*, *Maf*, or both *Prdm1* and *Maf*. The dot size represents the percentage of cells per cell cluster expressing the gene in question and the expression level indicated by the colour-scale.

### Genes expressed in human IBD are controlled by Blimp-1 and c-Maf

RNA-seq data from colon biopsies from published studies of IBD, including Crohn’s Disease (CD) and Ulcerative Colitis (UC), were obtained from the GEO (GSE193677; GSE126124)^38, 39^. Modular analysis was performed to display clusters of up- or down-regulated genes as compared to controls and modules of interest were selected and manually annotated (Fig. 8; Extended Supplementary Data Tables 7). The genes from the human ‘Immune effector’, and the ‘Granulocyte,Myeloid’ modules, which were more highly expressed in adult UC which usually manifests as inflammation of the colon, were selected and used to interrogate our mouse LPL scRNA-Seq data set from the *H. hepaticus* infected mice (Fig. 8a,b,c). Genes within the human IBD ‘Immune effector’ module were found to be largely expressed in the ‘B cells 1’, ‘Macrophage/Myeloid leukocyte’, ‘T cells Ctla4 high’ and ‘T cells Icos high’ in the mouse LPL scRNA-Seq clusters from infected *Prdm1*^fl/f^*Cd4*^Cre^ mice and to a greater extent the *Maf*^fl/fl^*Cd4*^Cre^ and double deficient *Prdm1*^fl/f^*Maf*^fl/f^*Cd4*^Cre^ mice (Fig. 8d). Expression of genes from the human ‘Granulocyte,Myeloid’ module, was largely confined to the mouse scRNA-Seq ‘Epithelial’ and ‘Macrophage/Myeloid leukocyte’ clusters, and expressed at the highest level in the *Maf*^fl/fl^*Cd4*^Cre^ and double deficient *Prdm1*^fl/f^*Maf*^fl/f^*Cd4*^Cre^ mice (Fig. 8e). T cell/ILC related genes shown earlier (Fig. 3b,d, k,l,n; Fig. 8g) to be expressed at the highest level in the LPL from *H. hepaticus* double deficient *Prdm1*^fl/f^*Maf*^fl/f^*Cd4*^Cre^ infected mice were found to be contained within these human IBD modules and their expression, including that of *IFNG, CSF2, IL18R1, IL- 12RB1, IL12RB2, IL23R* was found to be increased in CD and to a greater extent UC as compared to healthy controls (Fig. 8f,g; Extended Data Supplementary Table 7). Additionally, identified within the human IBD “Immune effector module” IL21 was found to be over-expressed in human IBD biopsies, particularly in UC, and showed the highest abundance in the LPLs from *H. hepaticus* double deficient *Prdm1*^fl/f^*Maf*^fl/f^*Cd4*^Cre^ infected mice (Fig. 8f,g), suggesting that in this case IL-21 is associated with a type I response. Other T cell/ILC associated genes, most highly expressed in LPL from infected *Maf*^fl/fl^*Cd4*^Cre^ mice (Fig. 3c,n), including *IL17A,* and now also *IL17F,* were highly expressed in human IBD, again to a higher extent in UC, as compared to controls (Fig. 8f,g; Extended Data Supplementary Table 7). Multiple genes associated with myeloid cells and innate immunity, which we showed earlier were most highly expressed in the LPL from infected *Maf*^fl/fl^*Cd4*^Cre^ mice (Fig. 5h; Fig. 8g), were over-expressed in human IBD, including *IL1B, IL1A, CSF3, ACOD1, OSM, S100A8, S100A9, CXCR2* (Fig. 8f). *ALPK2* now shown to be over-expressed in human IBD, mostly in UC, was however equally expressed in *Maf*^fl/fl^*Cd4*^Cre^ and double deficient *Prdm1*^fl/f^*Maf*^fl/f^*Cd4*^Cre^ mice (Fig. 8f,g), while *CXCL10* was increased in both UC and CD, and highest in the double deficient LPL (Fig. 8f,g). Other genes identified within the human IBD ‘Granulocyte,Myeloid’ module, including *NOD2, FCGR3* and *PROK2*, were most highly expressed in the biopsies from UC (Fig. 8f) and we show here that these genes are most highly abundant in the *Maf*^fl/fl^*Cd4*^Cre^ mice (Fig. 8g). Our findings indicate that genes which show increased expression in human IBD colon biopsies, including those with mutations which have been linked to an increased risk of CD and/or UC, such as *NOD2, IL23R, IL21* and *IFNG*^11, 40, 41^ are differentially abundant in the LPL from *H. hepaticus* infected mice with T cell-specific deletion of *Prdm1, Maf* or both transcription factors, suggesting that these mouse models may reflect different pathobiological mechanisms relevant in human IBD.

## DISCUSSION

Intestinal immune responses are tightly controlled to tolerate commensal microbiota whilst allowing control of invading pathogens. IL-10 production by CD4^+^ T cells is critical in regulating immune responses to avoid intestinal immune pathology in response to any potentially disease-causing microorganisms, which under normal circumstances do not cause adverse effects. The transcription factors Blimp-1, encoded by *Prdm1,* and c-Maf are co-dominant regulators of *Il10* in CD4^+^ T cells. We show herein that mice with T cell-specific deletion of *Prdm1, Maf* or the combination of both transcription factors do not develop inflammatory intestinal pathologies at the steady state but only upon infection with the pathobiont *H. hepaticus*. Using combined bulk tissue and single-cell RNA-Sequencing of LPL, together with ATAC-Sequencing and RNA-Seq of LPL purified CD4+ T cells, we show that during oral infection with *H. hepaticus* Blimp-1 and c-Maf both positively regulate *Il10* gene expression, but also cooperate and yet differentially negatively regulate a large network of proinflammatory effector responses. LPLs from the infected double deficient *Prdm1*^fl/fl^*Maf*^fl/fl^*Cd4*^Cre^ mice showed a high-level Type I immune response with increased TH1, IFN-γ^+^GM-CSF^+^effector cells and increased *Ifng, Csf2, Il23r*, *ll12rb1, Il12rb2* expression. This Type I response was also increased, but to a lesser extent, in the LPLs from infected *Prdm1*^fl/fl^*Cd4*^Cre^ and *Maf*^fl/fl^*Cd4*^Cre^ mice single T cell-specific transcription factor deleted mice. On the other hand, c-Maf negatively regulated a Type 17 response, with increased *Il17a* expression and a pronounced signature of neutrophils, myeloid cells and innate immunity in the LPLs from the *H. hepaticus* infected *Maf*^fl/fl^*Cd4*^Cre^ mice, which was less marked in the double *Prdm1*^fl/fl^*Maf*^fl/fl^*Cd4*^Cre^ LPLs. Thus, Blimp-1 and c-Maf cooperate to control common and distinct gene networks in T cells by distinct direct and shared actions on proinflammatory cytokines, over-and-above their direct stimulatory effects on *Il10,* subsequently controlling different effector immune responses and intestinal pathologies. Genes over-expressed in human IBD showed differential expression in the LPL from infected mice in the absence of T cell-specific expression of *Prdm1* or *Maf,* potentially revealing T cell regulated mechanisms relevant to human disease.

Our findings that T cell-specific deletion of either *Prdm1*, *Maf* or both transcription factors did not result in overt inflammation of the colon in the steady state are in keeping with most reports that *Maf^fl/fl^Cd4*^Cre^ or *Maf^f^*^l/fl^*Foxp3*^Cre^ mice do not develop colitis^13, 14, 16, 17^ in the absence of challenge, although there are a few conflicting reports of spontaneous mild colitis, but only in a small percentage of aged mice with either a Treg or T-cell-specific deletion in *Maf*^17, 18^. Likewise, there are conflicting reports in T-cell-specific *Prdm1*-deficient mice, with some indicating spontaneous colitis^22, 23, 26, 42^ while in other reports no signs of colitis were observed^24, 25^, the latter in keeping with our findings. One very recent study reported spontaneous colitis in double *Prdm1*^fl/fl^*Maf*^fl/fl^*Cd4*^Cre^ mice^14^, in contrast to our findings in the same mice, likely reflecting the controlled microbiota in our *vivarium*. It is thus likely that reports of spontaneous colitis in mice with a T cell-specific deletion of *Prdm1, Maf* or both transcription factors, which has been associated with either Tregs losing their immunosuppressive function^14^ or increased frequencies of TH17 cells^26^, are likely triggered by undefined microbiota or by undiagnosed infection with pathobionts. Microbiota or pathobionts may be maintaining TH17 cells at the steady state as has been described^16, 43, 44^ and very recently shown to be under c-Maf-dependent Treg control^16^. Our findings that mice with T cell-specific deletion of *Prdm1, Maf* or both transcription factors do not exhibit any signs of intestinal inflammation, provided us with an excellent baseline to determine the role of these transcription factors in regulation of the intestinal immune response to a defined pathobiont. We now show that infection with the pathobiont *H. hepaticus* led to graded increases in colitis, with the *Prdm1*^fl/fl^*Cd4*^Cre^ showing mild colitis, the *Maf*^fl/fl^*Cd4*^Cre^ showing moderate colitis and the double deficient *Prdm1*^fl/fl^*Maf*^fl/fl^*Cd4*^Cre^ showing severe colitis. No intestinal pathology was observed in the control *H. hepaticus* infected mice unless they were co-administered anti-IL-10R mAb in keeping with previous reports^8, 45, 46, 47, 48^. Since no increase in pathology was observed in the *Prdm1*^fl/fl^*Maf*^fl/fl^*Cd4*^Cre^ mice *H. hepaticus* infected mice when co-administered anti-IL-10R mAb, this suggested that the high level of intestinal pathology resulted from effects of both *Prdm1* and *Maf* on other immune factors, in addition to their co-dominant role in *Il10* gene regulation.

The distinct cellular and immune landscapes in the T cell-specific transcription factor deficient mice during *H. hepaticus* infection may represent distinct pathobiologic mechanisms of human IBD. This is supported by our findings that T cell/ILC associated genes such as *IFNG, IL12RB2, IL23R*, *IL18R, CSF2, IL17A* and *IL17F* that were highly expressed in colon biopsies from human IBD, as previously reported ^11, 38, 39, 40^, showed differential expression in the LPL from double deficient *Prdm1*^fl/fl^*Maf*^fl/fl^*Cd4*^Cre^ and *Maf*^fl/fl^*Cd4*^Cre^ mice and enriched in Foxp3^-^ effector T cells and ILC as shown by scRNA-seq data. Many of these proinflammatory genes have been reported as susceptibility loci for IBD and have highlighted shared genetic risk across populations^11, 40, 49^, in addition to mutations in the IL-10R^10^. Moreover, as discussed earlier, *Prdm1* has been reported to harbour rare missense mutations in PR domain-containing 1 (PRDM1) associated with IBD and these mutations resulted in increased T cell proliferation and production of proinflammatory cytokines, such as IFN-γ, upon activation^12^. Moreover, the Notch/STAT3-Blimp-1/c-Maf axis, shown to be a common anti-inflammatory pathway in T cells has been shown to be defective in in effector CD4+ T cells from Crohn’s disease patients^50^. While both Blimp-1 and c-Maf positively and directly regulate *Il10* gene expression as previously reported^3, 4, 5, 13, 14, 51^, here we show that they also directly control *Ifng, Il17a* and *Csf2* expression, as supported by common and distinct binding sites coinciding with open chromatin (ATAC-seq) sites from sorted CD4^+^ T cells from *H. hepaticus* infected mice. This is supported by our findings that blocking IL-10 signalling during *H. hepaticus* infection did not lead to enhanced pathology in the double deficient transcription factor deficient mice, reinforcing that the action mechanisms of Blimp-1 and c-Maf during immunoregulation of colitis are beyond IL-10 regulation.

The enhanced Type 1 response accompanying the intestinal pathology in the *Prdm1*^fl/fl^*Maf*^fl/fl^*Cd4*^Cre^ *H. hepaticus* infected mice and to a lesser extent the single T cell-specific transcription factor deficient mice, is in keeping with earlier studies implicating TH1 cells and IFN-γ in colitis^52^. Moreover, the potential role of TH1, IFN-γ^+^GM-CSF^+^ effector cells in the exacerbated intestinal pathology observed in the LPLs from infected *Prdm1*^fl/fl^*Maf*^fl/fl^*Cd4*^Cre^ is supported by previous reports showing that GM-CSF drives IL-23-driven colitis during infection of mice with *H. hepaticus* combined with anti-IL-10R treatment^53^. Although IL-23 receptor signalling has been strongly linked with induction of a TH17 response, our findings show increased expression of the *Il23r* by bulk tissue RNA-seq and scRNA-seq in LPL from *Prdm1*^fl/fl^*Cd4*^Cre^ and maximally in the *Prdm1*^fl/fl^*Maf*^fl/fl^*Cd4*^Cre^ mice, accompanying high levels of *Ifng* and *Csf2* is in keeping with reported roles for IL-23 in promoting IFN-γ production and intestinal inflammation^46, 54^. The significant increase in *Il17a* expression in the LPL from *H. hepaticus* infected *Maf*^fl/fl^*Cd4*^Cre^ mice supported a major role for c-Maf in negatively regulating IL-17 responses, in keeping with a previous report *in vitro* highlighting c-Maf as a repressor of TH17 responses^15^. However, since RNA-Seq from sorted CD4^+^ T cells and scRNA-Seq analysis of the ‘T cells Ctla4 high*’* cluster from LPL from *H. hepaticus* infected control mice showed a similar level of expression to that of *Il17a* in uninfected mice, this suggests that microbiota may be maintaining TH17 cells at the steady state as has been described^16, 43, 44^. The source of the increased *Il17a* in the LPL from *H. hepaticus Maf*^fl/fl^*Cd4*^Cre^ mice above that of infected control mice may be attributed to the increased abundance of TH17 cells, or to other IL-17A producers such as ILC3 or γ8 T cells as recently suggested^45^..

The total increase in the IL-17 response in the LPL from *Maf*^fl/fl^*Cd4*^Cre^ mice may explain the increased numbers of neutrophils observed by flow cytometry and scRNA-seq, in keeping with many report showing that IL-17 promotes neutrophil recruitment and function during many infections^55^. A role for neutrophils in the intestinal pathology arising in infected *Maf*^fl/fl^*Cd4*^Cre^ mice is in keeping with their association with human IBD and may reveal pathways for pathobiologic mechanisms in keeping with reports of IL-1-driven stromal-neutrophil interactions reported to define a subset of patients with IBD that does not respond to therapies^33^. Other neutrophil/myeloid associated genes, *PROK2* (prokineticin 2) and *FCGR3*, which were identified in the human IBD granulocyte/myeloid modules and were most highly expressed in the biopsies from UC patients, were most abundant in the LPLs from *H. hepaticus* infected *Maf*^fl/fl^*Cd4*^Cre^ mice, potentially underpinning pathobiologic mechanisms for human IBD. The greatest increase in neutrophil numbers and neutrophil associated genes was observed in the *Maf*^fl/fl^*Cd4*^Cre^ infected mice and to a lesser extent in the double deficient *Prdm1*^fl/fl^*Maf*^fl/fl^*Cd4*^Cre^ infected mice. This may suggest cross regulation of the IL-17 responses by IFN-γ^56^, which would result indirectly in reduced neutrophil responses. Alternatively, neutrophil migration or function by genes elevated in the absence of *Prdm1* in T cells in the double deficient *Prdm1*^fl/fl^*Maf*^fl/fl^*Cd4*^Cre^ infected mice, such as IFN-γ, may account for reduced neutrophil activity, in keeping with reports that IFN-γ restrains neutrophil function during infection^57^.

Inference of putative cell-to-cell communication networks using scRNA-seq data, revealed a graded increase in interactions between a cluster of T cells and myeloid cells in the *H. hepaticus* infected *Prdm1*^fl/fl^*Cd4*^Cre^, *Maf*^fl/fl^*Cd4*^Cre^ and double deficient *Prdm1*^fl/fl^*Maf*^fl/fl^*Cd4*^Cre^ mice as compared to the control infected mice, and this was recapitulated by immunofluorescent staining of colon sections. This increase in T cell-myeloid cell interactions might also be attributable to an increase in chemokine and cytokine ligand-receptor interactions. In line with these results, a cluster of genes associated with innate immunity which included cytokines previously associated with IBD, such as *Il1a*^32^, *Osm*^11, 58^ and other myeloid associated genes, *e.g. Acod1*, was elevated in the LPL from *H. hepaticus* infected *Maf*^fl/fl^*Cd4*^Cre^ and to a lesser extent double deficient *Prdm1*^fl/fl^*Maf*^fl/fl^*Cd4*^Cre^ infected mice as detected by both bulk tissue RNA-seq and scRNA-seq.

While our scRNA-Seq data revealed that T cell-derived *Il10 was* expressed mostly by Foxp3^+^ Treg, expression of *Ifng, Il17a* and *Csf2* was largely expressed by *Foxp3*^-^ T effector cells. Our findings support previous reports that Foxp3^+^ Treg are the major producers of *Il10* during *H. hepaticus* infection. However, although Foxp3^+^RORgt^+^ Treg were almost completely abolished in the LPL from infected *Maf*^fl/fl^*Cd4*^Cre^ mice in keeping with a previous report^31^, the LPL from the infected double deficient *Prdm1^fl^*^/fl^*Maf*^fl/^fl*Cd4*^Cre^ mice expressed the lowest levels of *Il10* and exhibited the maximum pathology indicating that Foxp3^+^RORgt^+^ T cells are not the only IL-10 producing Foxp3^+^ Treg population regulating intestinal pathology. Intestinal Tregs have been recently referred to as Foxp3^+^ Treg effector cells which exert their effects in lymphoid aggregates in the LP, since they show increased expression of *Areg, Gzmb, Icos, Tigit, Tnfrsf4* (OX40), and *Tnfrsf18* (GITR), in addition to IL-10^45^. Whilst our findings support the increased expression of these effector genes in the Foxp3^+^ Tregs, we show that these genes were also increased in Foxp3^-^ CD4^+^ T cells, in the LPL from *H. hepaticus* infected *Prdm1^fl/fl^Cd4^Cre^*, *Maf*^fl/fl^*Cd4*^Cre^ and double deficient *Prdm1^fl^*^/fl^*Maf*^fl/fl^*Cd4*^Cre^ mice as compared to controls, although to a much greater extent in the Foxp3- T cells (data not shown). Our findings are in keeping with previous reports that c-Maf controls Treg cell-derived IL-10 production and controls intestinal TH17 responses^16, 17^. However, our findings that c-Maf-deficient T cells that are Foxp3^-^ express the maximal levels of *Il17* in the LPLs from infected *Maf*^fl/fl^*Cd4*^Cre^ mice, suggest that c-Maf is controlling *Il17* expression in non-Treg IL-17-producing effector T cells also. This is supported the large increase in *Il17a* expression and *Il17a* expressing cells that we observe by bulk tissue and scRNA-seq of whole LPL in *H. hepaticus* infected mice *Maf*^fl/fl^*Cd4*^Cre^ mice over that of control infected or uninfected mice. The source of low levels of Blimp-1 and c-Maf-dependent *Il10* expression by Foxp3^-^ T cells, could potentially be TH cells or Tr1 cells^17, 59^, although the latter could have arisen by chronic stimulation of TH17 or TH1 cells as previously reported^60, 61^, or that Foxp3 expression has been lost on peripherally induced Treg cells^62^. Regardless, it is likely that Foxp3^+^ Treg are the major producers of *Il10* during *H. hepaticus* infection which control the intestinal inflammation. The CD4^+^ T cell source of IL-10 controlling immune responses to limit host damage is likely to be dictated by the immune response is to intestinal microbiota, pathobionts, different pathogens or different isolates of the same pathogen, and may possibly change at different stages of infection and/or anatomical locations as we previously discussed^4^.

Together our findings show that Blimp-1 and c-Maf co-operate to positively regulate *Il10* expression and to directly control genes encompassing the IL-23, IFN-γ, TH1 gene axis and a TH1 IFN-γ^+^GM-CSF^+^ effector cell response to prevent severe bacterial-induced colitis. Both transcription factors additionally indirectly regulated innate immune responses, although *Maf* uniquely regulated a pronounced signature of neutrophil activation, increased neutrophil numbers and increased *Il17a* expression. Thus, Blimp-1 and c-Maf control common and distinct gene networks that regulate qualitatively different pathologies, via positive regulation of *Il10* and importantly also by negative regulation of overlapping and distinct gene-networks in T cells, which subsequently control proinflammatory innate and CD4^+^ T cell effector pathways. That genes of the innate and adaptive immune response which were differentially expressed in the LPL of *H. hepaticus* infected mice deficient in T cell-specific *Prdm1* or *Maf* were elevated in gene sets from biopsies of patients with IBD^38, 39^ may provide novel pathobiologic mechanisms of human disease and additional targets for therapeutic development for IBD^63^. Our study highlights the power of applying genome-wide analyses of transcriptional regulation of effector immune responses using both bulk and scRNA-Seq and ATAC-Seq to reveal gene networks, validated by complementary flow cytometry analysis and immunofluorescent staining of colon biopsies, to reveal qualitative and quantitative distinct forms of pathobiont-induced colitis which may help delineate pathways of human disease.

## METHODS

### Animals

Mice were bred and maintained under specific pathogen free conditions in accordance with the Home Office UK Animals (Scientific Procedures) Act 1986. Age-matched male or female mice were used for experiments. *Maf*^fl/fl^ mice were provided by M. Sieweke and C. Birchmeier (Max Delbrück Centre for Molecular Medicine, Germany) ^64^ and backcrossed to C57BL/6J for ten generations and then crossed to *Cd4*^Cre^ mice to generate *Maf*^fl/fl^*Cd4*^Cre^ mice as described in ^13^. *Prdm1*^fl/fl^ mice were purchased from the Jackson Laboratory, backcrossed to C57BL/6J for four generations and then crossed to *Cd4*^Cre^ mice to generate *Prdm1*^fl/fl^*Cd4*^Cre^ mice. *Prdm1*^fl/fl^*Maf*^fl/fl^*Cd4*^Cre^ and *Prdm1*^fl/fl^*Maf*^fl/fl^ control mice were generated in-house by crossing *Maf*^fl/fl^*Cd4*^Cre^ with *Prdm1*^fl/fl^*Cd4*^Cre^ mice. All animal experiments were carried out in accordance with UK Home Office regulations and were approved by The Francis Crick Institute Ethical Review Panel.

### *H. hepaticus* colitis model and antibody treatment

*H. hepaticus* (NCl- Frederick isolate 1A, strain 51449) was grown under anaerobic gas conditions 10% CO2, 10% H2/N2 (BOC) for 3 days on blood agar plates containing 7% laked horse blood (Thermo Scientific) and the *Campylobacter* selective supplement “Skirrow” containing the antibiotics trimethoprim, vancomycin and polymyxin B (all from Oxoid). Bacteria were harvested and then transferred to and expanded, again under the anaerobic gas conditions above, for 3-4 days to OD 0.6 in tryptone soya broth (Oxoid) supplemented with 10% FCS (Gibco) and the antibiotics above. For infection, mice received 1x10^8^ colony forming units (c.f.u.) of *H. hepaticus* by oral gavage using a 22G curved blunted needle on Day 0 and Day 1. Uninfected mice were housed the same animal facility and only received antibody treatment. For experiments with anti-IL-10R: 1mg of either anti-mouse IL-10R (CD210) [Clone 1B1.3A, Rat IgG1, kappa] blocking antibody, or Rat IgG1 [Clone GL113, Rat IgG1] isotype control was administered on Day 0 and Day 7.

### Histopathology assessment

To assess the severity of colitis in *H. hepaticus* infected mice, in addition to uninfected and steady state aged mice (age 24-30 weeks), formalin-fixed paraffin-embedded cross-sections of proximal, middle and distal colon were stained with H&E and scored by two board-certified veterinary pathologists on a scale of 0-3 across four parameters to give a maximum score of 12. The four parameters included: epithelial hyperplasia and/or goblet cell depletion, leucocyte infiltration into the lamina propria, area affected, and markers of severe inflammation, which included crypt abscess formation, submucosal leucocyte infiltration, crypt branching, ulceration and fibrosis. Representative images of H&E-stained colon sections were then taken by the pathologists using a light microscope and a digital camera (Olympus BX43 and SC50).

### Immunostaining of colon

Proximal, middle and distal colon formalin-fixed paraffin-embedded cross-sections were de-waxed and re-hydrated before performing automated staining on the Leica BOND Rx Automated Research Stainer. Samples were treated with BOND Epitope Retrieval Solution 1 (Leica AR9961) for CD4 and CD68 antibodies, and BOND Epitope Retrieval Solution 2 (Leica AR9640) for MPO antibody. Permeabilization was performed in 3% hydrogen peroxide solution (hydrogen peroxide Fisher chemical code H/1750/15) and incubated in 1% BSA blocking buffer (BSA Sigma Aldrich A2153-100G, 1003353538 source SLBX0288). A multiplex panel of antibodies against CD4 (Rabbit, Abcam ab183685, clone EPR19514, 1:750 dilution), CD68 (Rabbit, Abcam ab283654, clone EPR23917-164, 1:2500 dilution) and MPO (Goat, R&D Bio-Techne AF3667, 1:200 dilution). Leica Novolink Polymer (anti-Rabbit, RE7161) was used as a secondary detection for primary antibodies raised in rabbit (CD4 and CD68) and Horse anti-goat IgG polymer reagent (Impress HRP)(Vector Laboratories 30036) for primary antibody MPO raised in goat. Samples were then incubated with Opal 690 (Akoya OP-001006) for CD4, Opal 520 (Akoya OP-001001) for CD68 and Opal 570 (Akoya OP-001003) for MPO, followed by DAPI nuclear counterstain (Thermo Scientific 62248, 1:2500 dilution). Slides were scanned using Akoya’s PhenoImager HT at 20x and viewed in Akoya inForm Automated Image Analysis Software. Scanned slides were imported into QuPath (version 0.4.3) for image analysis. Cell segmentation was performed using the Stardist extension on the DAPI channel in QuPath. Machine learning was used to train object classifiers on representative regions of each experimental group for CD4, CD68 and MPO markers. Exported data was used to determine the number of positive cells per area (um^2^) in gut sections of mice.

### Isolation of colon lamina propria leukocytes

LPLs were isolated from 1.0-1.5cm pieces of the proximal, middle and distal colon from individual mice, which were cleaned to remove faeces, opened up lengthwise and harvested into Dulbecco’s PBS with no Ca^2+^ and Mg^2+^ ions (Gibco) containing 0.1% (v/v) Bovine Serum Albumin Fraction V (Roche) (PBS+BSA). To remove the epithelium and intraepithelial lymphocytes, colonic tissue was incubated for 40 min at 37°C with 220rpm shake in 10ml of RPMI (Lonza, BE12-702F) supplemented with 5% (v/v) heat-inactivated FCS and 5mM EDTA (RPMI+EDTA). A second RPMI+EDTA wash was performed for 10 min, after which the tissue left standing at room temperature in 10ml RPMI (Lonza, BE12-702F) supplemented with 5% (v/v) heat-inactivated FCS and 15mM HEPES (RPMI+HEPES) to neutralise the EDTA. Tissue was then digested at 37°C with 220rpm shake for 45 min in 10ml of RPMI+HEPES with 120µL of Collagenase VIII added at 50mg/ml in PBS (Sigma). The 10ml of digested tissue was then filtered through a 70µM filter into tube containing 10ml of ice-cold RPMI+EDTA to neutralise the Collagenase VIII and the cells centrifuged (1300rpm, 7 min, 4°C). The resulting pellet was then resuspended in 4ml of 37.5% Percoll (GE healthcare), diluted in PBS+BSA from osmotically normalised stock and centrifuged (1800rpm, 5 min, 4°C). After centrifugation the pellet was recovered, resuspended in conditioned RPMI and used for subsequent analysis by flow cytometry and RNA and DNA extractions.

### Flow cytometry of colon lamina propria leukocytes

For the analysis of intracellular cytokine expression, isolated colon LPLs from individual mice were transferred to 48- well plates and restimulated with conditioned RPMI media containing 500ng/ml Ionomycin (Calibiochem) and 50ng/ml Phorbol 12-myristate 13-acetate (Sigma) for 2 hours, after which 10µg/ml Brefeldin A (Sigma) was added to each well and the cells incubated for another 2 hours. All incubations were conducted at 37°C in a humidified incubator with 5% carbon dioxide. Following re-stimulation LPLs were harvested into cold Dulbecco’s PBS with no Ca^2+^ or Mg^2+^ ions (Gibco). LPLs were first Fc-blocked for 15 min at 4°C (24G2, Harlan) and then stained with extracellular antibodies: CD90.2 (53-2.1, PE, Invitrogen), CD4 (RM4-5, BV785, Biolegend), TCR-β (H57-597, APC-e780, Invitrogen), CD8 (53-6.7, BV605, Biolegend), and the UV LIVE/DEAD™ Fixable Blue dead cell stain (Invitrogen). LPLs were then fixed for 15 min at room temperature with 2% (v/v) formaldehyde (Sigma) and permeabilised for 30 min at 4°C, using permeabilization buffer (eBioscience) and stained with the following cytokine antibodies for 30 min at 4°C: IL-17A (eBio17B7, FITC, Invitrogen), IFN-γ (XMG1.2, PE- Cy7, BD), IL-10 (JES5-16E3, APC, Invitrogen), and GM-CSF (MP1-22E9, BV421, BD). For transcription factor expression analysis, isolated LPLs remained unstimulated, were Fc-blocked and stained with the same extracellular antibodies, plus Ly6G (1A8, PE-Dazzle, Biolegend), CD11b (M1/70, eFluor450, Invitrogen), CD19 (1D3, BV711, BD Biosciences) and UV dead cell stain as for the restimulated LPLs, and fixed for 30 mins at 4°C using FoxP3/transcription factor staining kit (eBiosciences). Following permeabilization for 30 min at 4°C, using permeabilization buffer (eBioscience) LPLs were then stained with the following transcription factor antibodies for 30 min at 4°C: RORγt (Q31-378, AF647, BD) and Foxp3 (FJK-16s, FITC, Invitrogen). After staining, cells were resuspended in sort buffer (2% FBS in PBS + 2mM EDTA) and analysed on the Fortessa X20 (BD) flow cytometer. Acquired data was analysed using FlowJo v10, with compensation performed using single colour controls from the cells and AbC™ total compensation beads (Invitrogen). Flow cytometry plots were concatenated for visualization purposes as follows, each individual acquisition file was down-sampled to the lowest number of events per genotype, thus resulting in a final concatenated file with even representation of each individual mouse per group. For intracellular cytokine staining, plots in Extended Fig. 4d,e,f, are comprised of n=5 for *Prdm1*^fl/fl^*Maf* ^fl/fl^ and n=4 for *Prdm1*^fl/fl^ *Cd4*^Cre^, *Maf* ^fl/fl^*Cd4*^Cre^, and *Prdm1*^fl/fl^*Maf* ^fl/fl^*Cd4*^Cre^. The transcription factor staining plots in Extended Fig. 4m are comprised of n=5 for *Prdm1*^fl/fl^*Maf* ^fl/fl^, n=2 for *Prdm1*^fl/fl^ *Cd4*^Cre^, n=4 for *Maf* ^fl/fl^*Cd4*^Cre^, and n=5 for *Prdm1*^fl/fl^*Maf* ^fl/fl^*Cd4*^Cre^. The extracellular marker staining plots in Fig. 5f are comprised of n=5 for *Prdm1*^fl/fl^*Maf* ^fl/fl^, n=5 for *Prdm1*^fl/fl^ *Cd4*^Cre^, n=5 for *Maf* ^fl/fl^*Cd4*^Cre^, and n=4-5 for *Prdm1*^fl/fl^*Maf* ^fl/fl^*Cd4*^Cre^ in each uninfected and infected group.

### Sorting by flow cytometry of CD4+ T cells from colon lamina propria

Colon LPLs were isolated as described earlier from individual mice and the cells harvested into cold Dulbecco’s PBS (no Ca^2+^/Mg^2+^ ions) (Gibco). Prior to the sort, colon LPLs from individual mice within some experimental groups were equally pooled as follows to allow for the sorting of n=3-6 biological replicates per experiment. In the uninfected groups for *Prdm1*^fl/fl^*Maf* ^fl/fl^ and *Prdm1*^fl/fl^*Maf* ^fl/fl^*Cd4*^Cre^ (n=12 mice were pooled to 3 give biological replicates). For the infected *Prdm1*^fl/fl^*Maf* ^fl/fl^ group (n=16 mice were pooled to 4 give biological replicates). Whereas for the following infected groups, individual mice were not pooled to give *Prdm1*^fl/fl^*Maf* ^fl/fl^*Cd4*^Cre^ (n=6), *Prdm1*^fl/fl^ *Cd4*^Cre^ (n=4) and *Maf* ^fl/fl^*Cd4*^Cre^ (n=6). For the FACS staining LPLs were first Fc-blocked for 15 min at 4°C (24G2, Harlan) and then stained with the extracellular antibodies: CD90.2 (53-2.1, PE, Invitrogen), CD4 (RM4-5, BV785, Biolegend), TCR-β (H57-597, APC-eFluor 780, Invitrogen), CD8 (53-6.7, BV605, Biolegend), and the UV LIVE/DEAD™ Fixable Blue dead cell stain (Invitrogen). Live CD4+ T cells (CD4+ TCR- β + CD90.2+ CD8-) were then sorted to over 95% purity on the FACS Aria III or FACS Aria Fusion cell sorters (both BD). Sorted cells were then used for subsequent RNA and DNA extractions.

### RNA-seq of colon lamina propria leukocytes

RNA was extracted from colon lamina propria leukocytes from individual mice using the QIAShredder and RNeasy Mini Kit with on-column DNase digestion, according to the manufacturer’s instructions (Qiagen). RNA-seq libraries were then made with total RNA using the KAPA RNA HyperPrep with RiboErase and unique multiplexing indexes, according to the manufacturer’s instructions (Roche). All libraries were then sequenced using the HiSeq 4000 system (Illumina) with paired-end read lengths of 100bp and at least 25 million reads per sample.

### scRNA-seq of colon lamina propria leucocytes

Isolated colon LPLs from individual mice (as detailed above) were filtered using a 70um filter cells and were suspended in PBS 0.04%BSA (UltraPure™ BSA; Invitrogen), for each sample an aliquot of cells was stained with AO/PI Cell Viability Kit (Logos Biosystems) and were counted with the LunaFx automatic cell counter. For all samples, cell viability prior to loading was >80%. As per the manufacturer’s instructions, the MasterMix was prepared as detailed in the Chromium Next GEM Single Cell 3’ Reagent Kit v3.1 (Dual Index) manual and 10,000 cells per sample were loaded into the 10x Chromium chips (10x Genomics). The 10x Chromium libraries were prepared and sequenced (paired-end reads) using the NovaSeq 6000 (Illumina).

### RNA-seq of sorted CD4+ T cells from colon lamina propria

RNA was extracted from flow sorted CD4+ T cells isolated from the colon lamina propria of individual mice using the QIAShredder and RNeasy Mini Kit with on-column DNase digestion, according to the manufacturer’s instructions (Qiagen). RNA-seq libraries were then made with total RNA using the NEBNext Single Cell/Low Input RNA Library Prep Kit for Illumina and unique multiplexing indexes, according to the manufacturer’s instruction (New England Biolabs). All libraries were then sequenced using the HiSeq 4000 system (Illumina) with paired-end read lengths of 100bp and at least 25 million reads per sample.

### ATAC-seq of sorted CD4+ T cells from colon lamina propria

ATAC-seq samples from isolated LPLs were prepared as outlined in^65^. For each sample, 50,000 cells were lysed in cold lysis buffer containing 10mM Tris-HCl, pH 7.4, 10mM NaCl, 3mM MgCl2, 0.1% Nonidet™ P40 substitute (all Sigma) and the nuclei incubated for 2 hours at 37°C with 50μl of TDE1/TD transposase reaction mix (Illumina). Tagmented DNA was then purified using the MinElute kit (Qiagen) and amplified under standard ATAC PCR conditions: 72°C for 5 min; 98°C for 30s and thermocycling at 98°C for 10s, 63 °C for 30s and 72°C for 1 min for 12 cycles. Each 50μl PCR reaction consisted of: 10μl Tagmented DNA, 10μl water, 25μl NEBNext High-Fidelity 2x PCR Master Mix (NEB), 2.5μl Nextera XT V2 i5 primer and 2.5μl Nextera XT V2 i7 primer (Illumina). NexteraXT V2 primers (Illumina) were used to allow larger scale multiplexing, these sequences were ordered directly from Sigma (0.2 scale, cartridge) and diluted to 100μM with 10mM Tris-EDTA buffer, pH8 (Sigma) and then to 25μM with DEPC-treated water (Ambion) for use in the reaction. Following amplification, ATAC-seq libraries were cleaned up using 90μl of AMPure XP beads (Beckman Coulter) and two 80% Ethanol washes whilst being placed on a magnetic plate stand, before being eluted in 1mM (0.1x) Tris-EDTA buffer, pH8 (Sigma) diluted with DEPC-treated water (Ambion). ATAC-seq libraries were then checked on the TapeStation/BioAnalyser (Agilent) before being sequenced on the HiSeq 4000 system (Illumina), with paired-end read lengths of 50bp and at least 50-80 million uniquely mapped reads per sample.

### Statistical analysis

All figure legends show the number of independent biological experiments performed for each analysis and replicates. Flow cytometry percentages and associated cell numbers were analysed as a one-way ANOVA with Tukey’s multiple comparisons test with 95% confidence interval for statistical analysis. All statistical analysis, apart from sequencing was carried out with Prism8 software (GraphPad, USA) (*=p≤0.05; **=p≤ 0.01; ***=p≤ 0.001, ****=p≤0.0001). Analyses for RNA-seq and ATAC-seq data were performed with R version 3.6.1 and Bioconductor version 3.9. Analyses for scRNA-seq data were performed with R version 4.1. and with Seurat version Seurat_4.1.1. Error bars and n values used are described in the figure legends.

### RNA-seq data processing and analysis

For bulk tissue LPL RNA-seq, adapters were trimmed using Skewer software version 0.2.2 ^66^ with the following parameters: “- m pe -q 26 -Q 28 -e -l 30 -L 100”, specifying the relevant adapter sequences. For sorted CD4+ T cell RNA-seq, adapters were trimmed using FLEXBAR software^67^, as recommended by NEB (the manufacturer) when using the NEBNext Single Cell/Low input RNA Library Prep Kit for Illumina. FLEXBAR was run following the provider’s suggested pipeline found in: https://github.com/nebiolabs/nebnext-single-cell-rna-seq. For both bulk tissue LPL RNA-seq and sorted CD4+ T cell RNA-seq reads were aligned to mm10 genome and the GENCODE reference transcriptome version M22 using STAR software version 2.7.1^68^, excluding multi-mapping reads by setting the parameter “outFilterMultimapNmax” to 1. In order to increase read mapping to novel junctions the parameter “twopassMode” was set to “Basic”. Raw gene counts were retrieved using QoRTs software version 1.1.8^69^. Normalised read counts were retrieved using DeSeq2 version 1.24.0^70^ and rlog transformed in order to visualise gene quantifications.

### Differential gene expression of bulk tissue LPL from *H. hepaticus* infected mice vs uninfected *Prdm1*^fl/fl^ *Maf*^fl/fl^ control

DeSeq2 ^70^ was used to obtain differentially expressed genes for each of the four *H. hepaticus* infected groups: *Prdm1*^fl/fl^*Maf* ^fl/fl^*, Prdm1*^fl/fl^*Cd4*^Cre^, *Maf* ^fl/fl^*Cd4*^Cre^, and *Prdm1*^fl/fl^*Maf* ^fl/fl^*Cd4*^Cre^ against the uninfected *Prdm1*^fl/fl^*Maf* ^fl/fl^ control. A gene was considered to be a statistically differentially expressed gene if the fold change >=1.5 and BH adjusted p value<0.05. Resulting in:

*Prdm1*^fl/fl^*Maf* ^fl/fl^ infected vs uninfected *Prdm1*^fl/fl^*Maf* ^fl/fl^: 5 DEG

*Prdm1*^fl/fl^*Cd4*^Cre^ vs uninfected *Prdm1*^fl/fl^*Maf* ^fl/fl^: 1207 DEG

*Maf* ^fl/fl^*Cd4*^Cre^ vs uninfected *Prdm1*^fl/fl^*Maf* ^fl/fl^: 1740 DEG

*Prdm1*^fl/fl^*Maf* ^fl/fl^*Cd4*^Cre^ vs uninfected *Prdm1*^fl/fl^*Maf* ^fl/fl^: 3392 DEG

### Cell enrichment and biological pathway annotation

The identified differentially expressed genes (in any condition) were subjected to *k-*means clustering using a *k*=9; the expression values for the DEG were standardised into z-scores and visualised in a heatmap (Fig. 1f). In order to provide a biological interpretation to these clusters, each cluster was subjected to “cell-type enrichment” and “biological pathways” annotation. The cell-type enrichment analysis used the cell-type signatures from^29^ and a Fisher’s exact test was performed to identify cell-type signatures enriched in each of the clusters, adjusted p-values were obtained using BH correction. The cell-type signatures that are statistically significant enriched (adjusted p-value<0.05) are shown in Fig. 1g. The R package “topGO” ^71^ was used to obtain the enriched biological processes in each cluster (Extended Fig. 1e). Additionally, an IPA “core analysis” (QIAGEN Redwood City, www.qiagen.com/ingenuity) was performed in order to identify the IPA pathways enriched in each cluster, the *Prdm1^fl/fl^Maf ^fl/fl^Cd4^Cre^*vs uninfected *Prdm1^fl/fl^Maf ^fl/fl^* expression values served as input for the calculation of the z-scores in the bar-plots depicted in Extended Fig. 1e, as this was the comparison that resulted in the highest amount of differentially expressed genes.

### Differential gene expression of sorted CD4+ T cells from *H. hepaticus* infected mice vs uninfected *Prdm1*^fl/fl^ *Maf*^fl/fl^ control

DeSeq2^70^ was used to obtain differentially expressed genes for the uninfected *Prdm1*^fl/fl^*Maf* ^fl/fl^*Cd4*^Cre^ and each of the four *H. hepaticus* infected groups: *Prdm1*^fl/fl^*Maf* ^fl/fl^*, Prdm1*^fl/fl^*Cd4*^Cre^, *Maf* ^fl/fl^*Cd4*^Cre^, and *Prdm1*^fl/fl^*Maf* ^fl/fl^*Cd4*^Cre^ against the uninfected *Prdm1*^fl/fl^*Maf* ^fl/fl^ control in Extended Fig. 4p. A gene was considered to be a statistically differentially expressed gene if the fold change >=1.5 and BH adjusted p value<0.05.

### ATAC-seq data processing and analysis

Paired end ATAC-seq reads from sorted CD4+ T cells, were quality controlled and adapters were trimmed using Skewer software version 0.2.2^66^ with the following parameters: “-m pe -q 26 -Q 30 -e -l 30 -L 50”, specifying “CTGTCTCTTATACAC” as reference adapter sequence to remove.

The reads were then aligned to mm10 genome using BWA-MEM ^72^, duplicate reads were removed with Picard ^73^ and SAMtools 1.3.1^74^ was used to discard discordant alignments and/or with low mapping qualities (mapQ<30). In order to account for transposase insertion, reads were shifted +4bp in the forward and -5bp in the reverse strand; moreover, read-pairs that spanned >99bp were excluded from further analyses as they would span nucleosomes^65^. MACS2 (version 2.1.1) was used to identify ATAC-seq peaks using the following parameters: “parameters --keep-dup all --nomodel -- shift -100 --extsize 200; q-value < 0.01”, in order to identify enrichment of Tn5 insertion sites^75^. DiffBind software version 2.0.2^76^ was used to generate raw counts underlying each ATAC-seq peaks. Furthermore, batch correction was performed on raw counts using the RUVSeq R package^77^ to remove batch effects that resulted from independent experiments. BeCorrect software^78^ was used to generate batch-corrected bigwig files, using the outputs from RUVseq software. The resulting batch corrected and normalised counts were used for visualisation. The R package “ggbio” was used to plot the genome browser tracks^79^.

### ChIP-seq data processing and analysis

Publicly available c-Maf ChIP-seq raw fastq files were obtained from GSE40918^15^ and Blimp-1 ChIP-seq raw fastq files were obtained from GSE79339^80^. Trimmomatic version 0.36 was used for quality control and trim adapter sequences using the following parameters: “HEADCROP:2 TRAILING:25 MINLEN:26”^81^. Trimmed reads were aligned to mouse genome mm10 with Bowtie 1.1.2 ^82^ with the parameters: “y -m2 --best --strata -S”. MACS2 2.1.1 was used with default parameters to identify ChIP-seq peaks, and peaks with a q-value<0.01 were defined as statistically significant binding sites. “bamCoverage” from DeepTools 2.4.2 was used to normalise ChIP-seq data to RPKMs and the R package “ggbio” was used to visualise the genome browser tracks^79^ together with the ATAC-seq data.

### IPA pathways

Th1 and Th17 pathways were constructed in and obtained from IPA signalling pathways library. Log2(Fold-changes) from the differential expression analyses outlined above were overlaid on the Th1 and Th17 pathways, all with a fixed scale of -5 (blue) to 3.5 (red).

### scRNA-seq data processing and analysis

Fastq files were aligned to the mm10 transcriptome, and count matrices were generated, filtering for GEM cell barcodes (excluding GEMs with free floating mRNA from lysed or dead cells) using Cell Ranger version 6.1.2. Count matrices were imported to R and processed using the Seurat library (version 4.0) following the standard pipeline ^83^. Low quality cells were removed, with cells kept for further analysis if they met the following criteria: the mitochondrial content was within 3 standard deviations from the median, more than 500 genes were detected and more than 1,000 RNA molecules were detected. DoubletFinder was used to identify doublets, assuming a theoretical doublet rate of 7.5% ^84^. All samples were integrated using the CCA method, implemented by Seurat’s functions FindIntegrationAnchors() and IntegrateData(), using the top 10000 variable features and the first 50 principal components. 23 clusters were identified from the integrated dataset, and were annotated with the scMCA R library (version 0.2.0) using the single cell Mouse Cell Atlas as reference ^30^ and the clustifyr R library using the Immgen reference dataset (version 1.5.1). A final manually curated annotation was assigned to clusters based on scMCA and clustifyr results, this resulted in the annotation of 17 distinct cell clusters. Marker genes for these cell clusters were identified using a Wilcoxon Rank Sum test, comparing each cluster to all other clusters, genes that were statistically significant (adjusted p value <0.05 and log2(FC)>0) were kept for further analysis.

### CellChat analysis

Cell-to-cell crosstalk was inferred using the R library CellChat version 1.1.3^37^ and the CellChat mouse database. The CellChat analysis was performed as outlined in the CellChat software manual, with the “population.size” parameter set to TRUE when computing the communication probability between clusters.

### Human IBD RNA expression data analysis

Publicly available human IBD RNA expression datasets were obtained from GSE193677 (adult; ^38^) and GSE126124 (paediatric; ^39^) were downloaded and RMA normalised using the limma package (version 3.50.0). From both datasets only colon biopsies from healthy controls and colon biopsies from untreated patients were used for further analysis. Normalised log2 expression values of the top 10,000 genes based on variances per dataset were used as input to WGNA (version 1.72.1) ^85^. Gene set modules were detected using a minimum module size of 30, and a deep.split of 2 for both datasets. This resulted in 12 modules for GSE193677 and 18 modules for GSE126124. GO biological process enriched in the modules were annotated using clusterProfiler (version 4.0.5). Using clusterProfiler results and manual curation a final annotation for some key modules was assigned. Furthermore, modules were tested using Qusage (version 2.26.0) using normalised log2 expression values as input for reciprocal datasets; statistically significant modules (adjusted p value<0.05) were plotted using ggcorrplot function in R. Genes within the modules derived from WGCNA were converted to mouse gene symbols using the bioMart (version 2.56.1) R library and genes not expressed in our mouse scRNA-seq dataset herein generated were filtered out. The remaining WGNA module genes were scored in our mouse scRNA-seq dataset using the AddModuleScore() function implemented in the Seurat R library.

### Data availability

The materials, data and any associated protocols that support the findings of this study are available from the corresponding author upon request. The RNA-seq datasets have been deposited in the NCBI Gene Expression Omnibus (GEO) database with the primary accession number GSE240422. Publicly available datasets used in this study include GSE40918 (c-Maf ChIP-seq), GSE79339 (Blimp-1 ChIP-seq), GSE193677 (adult IBD) and GSE126124 (paediatric IBD).

## ACKNOWLEDGMENTS

We wish to thank The Francis Crick Institute: Christine Graham (ex-AOG lab member) for training LSC in RNA library prep for RNA-Seq; Vicki Metzis for advice on ATAC-Seq; Vangelis Stavropoulos for lab support; Lucia Moreira-Teixeira for scientific input and help setting up and harvesting of *in vivo* experiments; Biological Services for breeding and maintenance of the mice used for experiments leading up to the current study, specifically, Anna Sullivan team (breeding); Experimental CL2 facility, Julie Bee and team and Jake Murphy and Peter Owers for oral gavage; Alina Janney Kennedy for helping the take-down of one *in vivo* experiment performed at the Kennedy Institute with LSC and CP; Advanced Sequencing Facility, Robert Goldstone and Maria Rodriguez for excellent project management of sequencing and Deb Jackson and Laura Cubitt for support of sequencing; Olga O’Neill, Equipment Park for QC; and Experimental Histopathology, Emma Nye, Mary Green, Ania Mikolajczak and team for their excellent work on multiplex immunofluorescent staining of colon sections and for their excellent support in preparing colon sections for histological analyses and Ellie Herbert for training of AOG lab members; Flow Cytometry members, including Andy Riddell, Phil Hobson and Sukhveer Purewal and team. We would like to thank would like to thank Hannah Painter, Crick AOG lab and now LSHTM for input to the manuscript, including downloading of human datasets from the GEO, careful review of all figures and suggestions for an improved publication and Kunyuan Tian for her input into and careful reading of the manuscript. At The Kennedy Institute we wish to thank the Powrie lab and Kennedy Institute BRF; and with help from Alina Janney and Elizabeth Mann who helped on the experimental harvest of *H. hepaticus* infected mice; and wish to thank Mathilde Pohin for her invaluable feedback on the manuscript and signatures. AOG would like to thank: Theodore Kapellos, Helmholtz Zentrum München-Deutsches Forschungszentrum für Gesundheit und Umwelt (GmbH), for excellent input to approaches to scRNA-Seq on neutrophils and bioinformatic analysis options; and Malte Lücken, Helmholtz Institute, Germany) for deep discussion and advice on bioinformatic analyses programmes for sc-RNA-Seq analyses for interrogating cell-cell interactions.

This whole study was funded by The Francis Crick Institute which receives its core funding from Cancer Research UK (FC001126), the UK Medical Research Council (FC001126), and the Wellcome Trust (FC001126); before that by the UK Medical Research Council (MRC U117565642). AOG, LC, MAM, VAS, XW, LMT, LG were supported by The Francis Crick Institute which receives its core funding from Cancer Research UK (FC001126), the UK Medical Research Council (FC001126), and the Wellcome Trust (FC001126); before that by the UK Medical Research Council (MRC U117565642); S.L.P. and A.S-B. were funded by the Royal Veterinary College and The Francis Crick Institute. WJB was supported by the AOG Wellcome Investigator Award. CP and FP were supported by a Wellcome Investigator Award to FP. Conflicts of interest: FP received grant funding or consultancy fees from Roche, Genentech, GSK, Janssen, Novartis and T Cypher.

## AUTHOR CONTRIBUTIONS

AOG, LSC and MAM co-designed the study, interpreted and analyzed the data and AOG and MAM co-wrote the paper with input from FP and WJB. All co-authors read and provided feedback for the paper. MAM, LSC, VAS, and WJB conducted *in vivo* experiments with support from XW and AAD at The Crick; LSC and CP, conducted the experiments at The Kennedy Institute: MAM, WJB, LSC and VAS analyzed the data and interpreted together with AOG; MAM analyzed the ATAC-Seq, ChIP-Seq, RNA-seq and scRNA-seq data; PC analyzed the scRNA-seq and human transcriptomic data obtained from GEO; HS performed the 10x Genomics reactions and libraries for the scRNA-seq; XW executed the genetics for obtaining and crossing all our T cell-specific transcription deficient mice and controls and designed, performed all screening and quality control and provided input throughout to the study; JB provided expert advice and input on the ATAC-Seq analysis and on the design of RNA-Seq and ATAC-Seq expts and feedback on the study and the paper; SP and ASB conducted all the colon pathology analyses on mice and provided all scoring and expert advice on the pathology data; AAD performed imaging, interpretation and analysis of the immunostained gut sections. AM developed, optimised and performed all the immunofluorescent staining for the gut sections and provided advice to AAD.

## Author information

Reprints and permissions information is available at www.nature.com/reprints. The author(s) declare no competing financial interests. Correspondence and requests for materials should be addressed to.

---

**Extended Fig.1.**
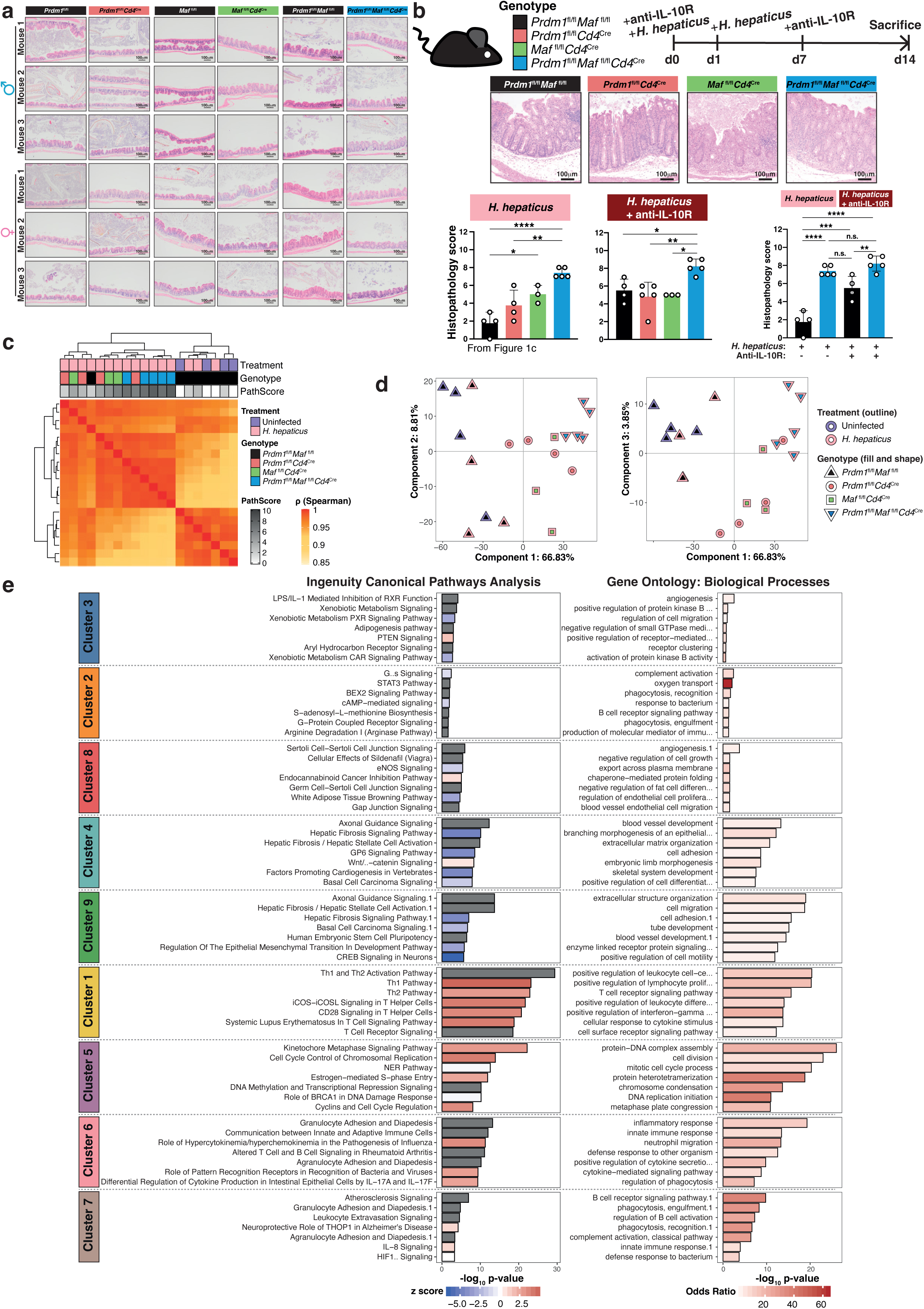
RNA-seq analysis of total colon LPL identifies changes in gene expression regulated by Blimp-1 and c-Maf upon *H. hepaticus* infection and highlights diverse biological processes and cell populations involved. **a)** Representative H&E colon sections from steady state mice aged to 24-30 weeks, scale bar = 100μm. Data representative of multiple female and male mice: *Prdm1*^fl/fl^ (n=8), *Prdm1*^fl/fl^*Cd4*^Cre^ (n=19), *Maf*^fl/fl^ (n=10), *Maf*^fl/fl^*Cd4*^Cre^ (n=22), *Prdm1*^fl/fl^*Maf*^fl/fl^ (n=36), *Prdm1*^fl/fl^*Maf*^fl/fl^*Cd4^C^*^re^ (n=38). **b)** Schematic of experimental method used to infect mice with *H. hepaticus* by oral gavage and treatment with anti-IL-10R blocking antibody. Representative H&E colon sections (scale bar = 100μm) from each genotype following infection with *H. hepaticus* and treatment with anti-IL-10R antibody and harvested on Day 14 with the corresponding colon histopathology scores (bottom panels). Each dot within the barplots represents an individual mouse analyzed. Graph shows mean ±s.d., analyzed by one-way ANOVA followed by Tukey’s post-hoc test (*=p value ≤ 0.05, **=p value ≤ 0.01, ***=p value ≤ 0.001, ****=p value ≤ 0.0001). **c)** Unsupervised hierarchical clustering of a pair-wise Spearman correlation and **d)** PCA plots showing on PC1 the separation of floxed control mice (uninfected and infected) versus *H. hepaticus* infected mice with *Cd4*^Cre^-mediated deletion of *Prdm1*, *Maf*, or *Prdm1* and *Maf*. **e)** Enriched IPA canonical pathways (left) and gene ontology biological processes (right) were obtained for the clusters of differentially expressed genes arising upon infection and *Cd4*^Cre^-mediated deletion of either *Prdm1*, *Maf*, or both *Prdm1* and *Maf* (related to Fig. 1f). Data from n=3-5 mice.

**Extended Fig.2.**
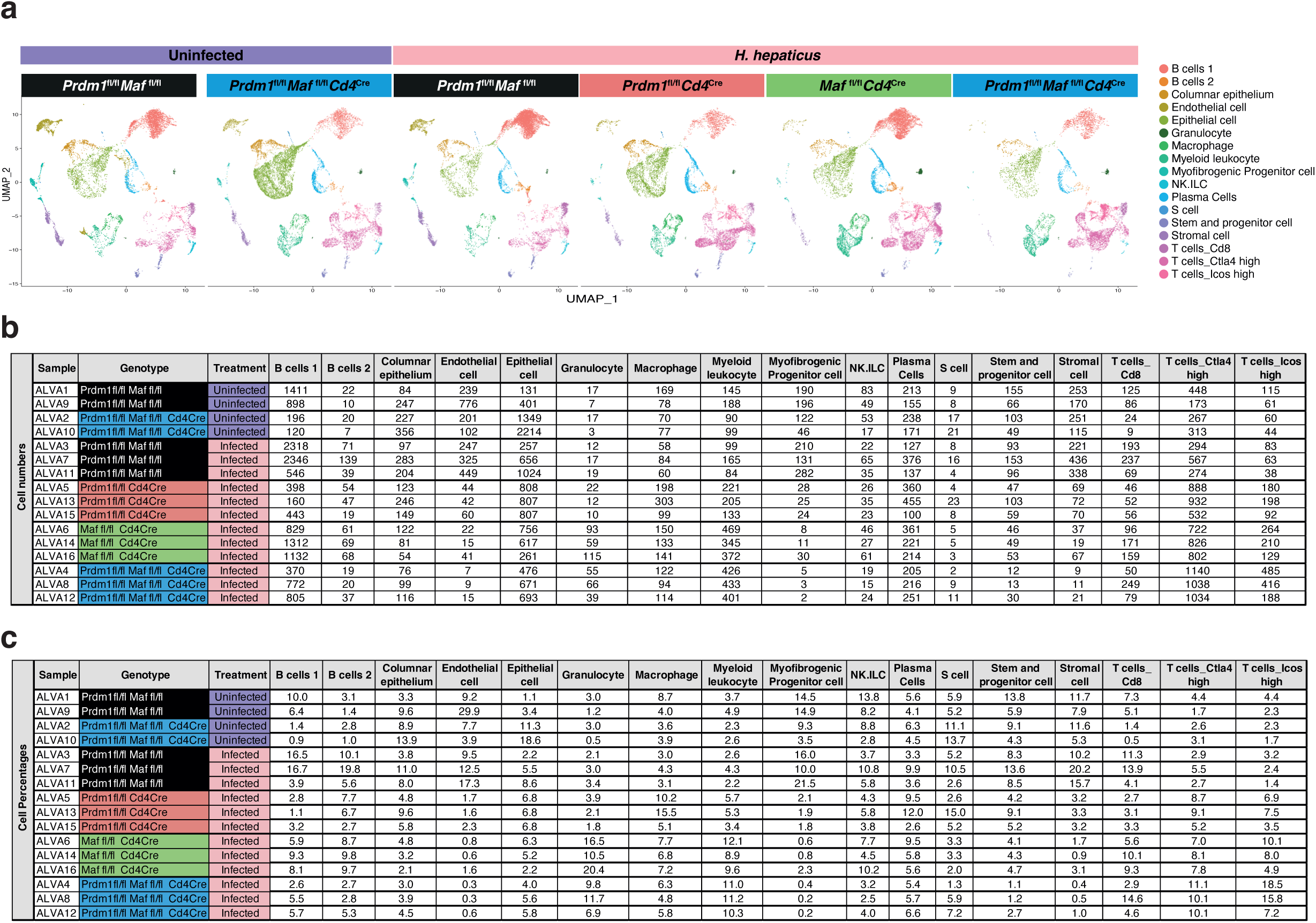
scRNA-seq analysis of total colon LPL identifies quantitative changes in cell populations affected by *Cd4*^Cre^-mediated deletion of *Prdm1*, *Maf*, or *Prdm1* and *Maf*. **a-c)** scRNA-seq was performed on total colon LPL isolated from uninfected *Prdm1*^fl/fl^*Maf*^fl/fl^ and *Prdm1*^fl/fl^*Maf*^fl/fl^*Cd4*^Cre^ control mice, and *H. hepaticus* infected *Prdm1*^fl/fl^*Maf*^fl/fl^ mice and mice with *Cd4*^Cre^-mediated deletion of either *Prdm1*, *Maf*, or both *Prdm1* and *Maf*. **a)** UMAP visualization of the integrated scRNA-seq dataset, plotted per condition and colored by assigned cell cluster. **b)** Total cell number and **c)** proportion of cells (%) successfully sequenced and passing quality control (Methods section) in each of the cell clusters identified. Data from n=2-3 mice.

**Extended Fig.3.**
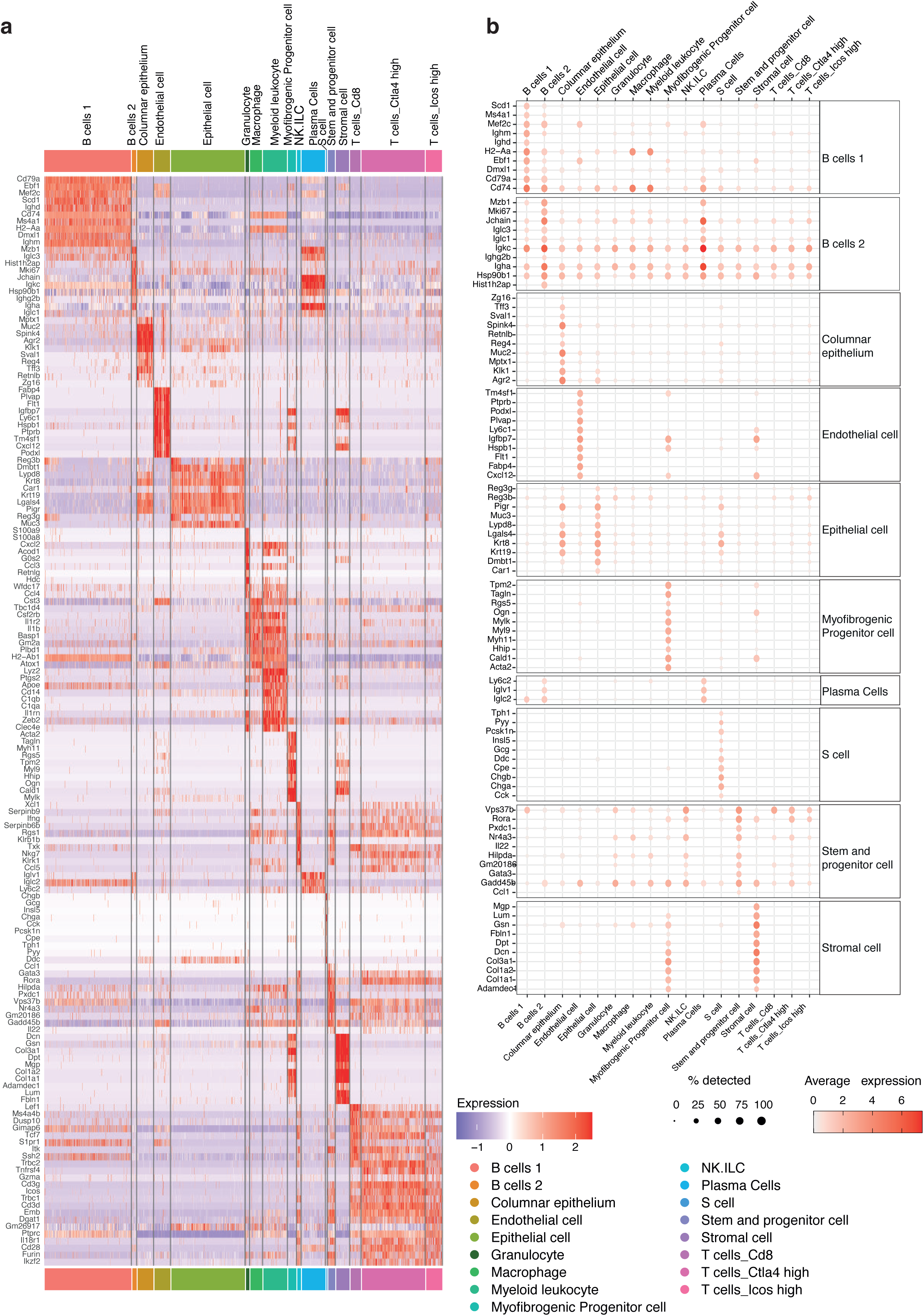
Marker genes identifying each of the 17 cell clusters in the scRNA-seq of total colon LPL from uninfected and *H. hepaticus* infected mice. **a)** Heatmap and **b)** dotplot (of cell clusters not in **Fig.2e**) of the top 10 differentially expressed marker genes of the identified cell clusters in our colon LPL scRNA-seq. Colored by the average gene expression across all cell clusters. In the dotplots, the dot size represents the percentage of cells per cell cluster expressing the gene in question.

**Extended Fig.4.**
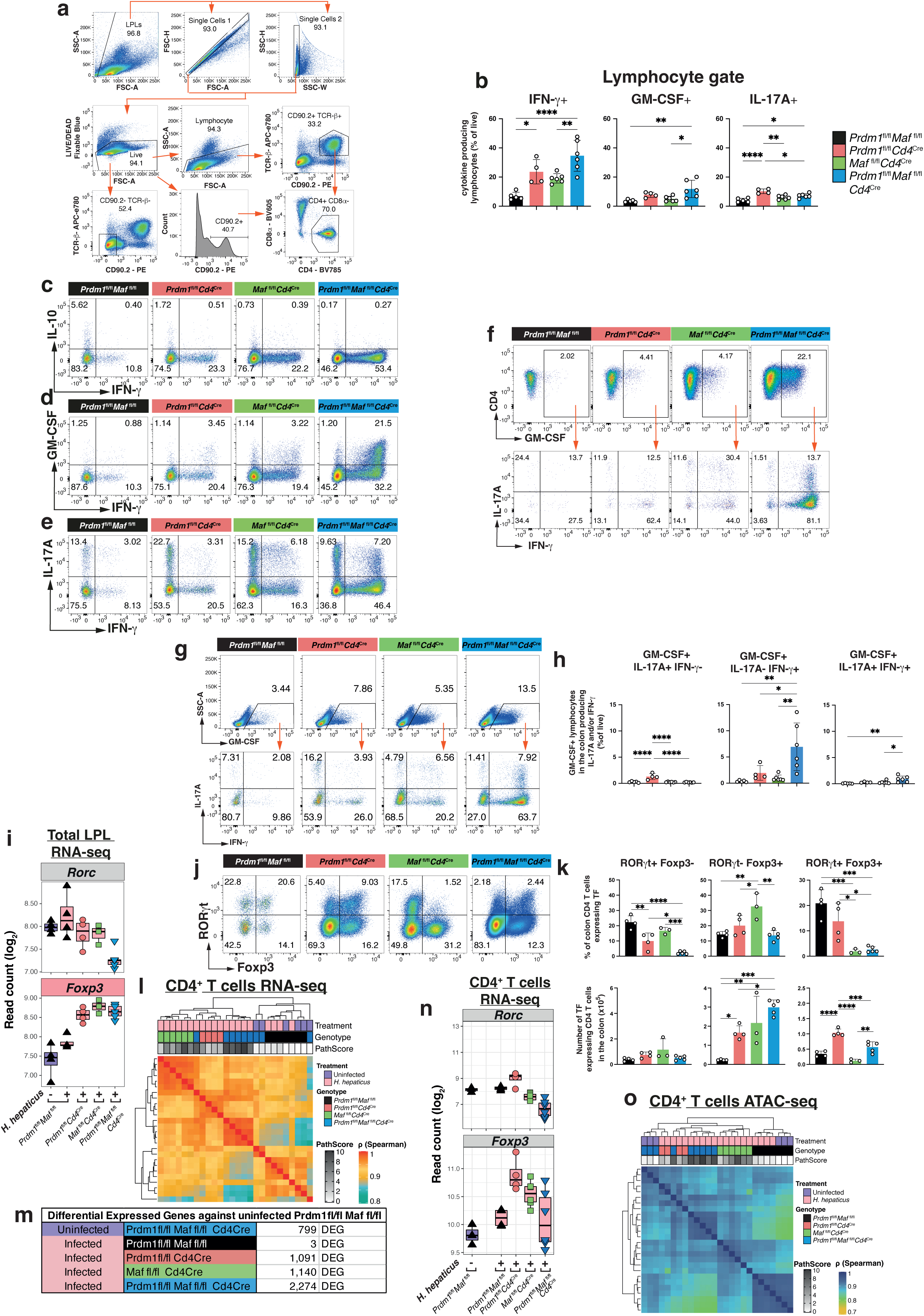
Flow cytometry analysis of colon LPL from *H. hepaticus* infected mice highlight increase pro-inflammatory cytokine production in *H. hepaticus* infected mice with *Cd4*^Cre^-mediated deletion of *Prdm1*, *Maf,* or both *Prdm1* and *Maf* when compared to infected *Prdm1*^fl/fl^*Maf*^fl/fl^ mice. **a)** Gating strategy for flow cytometry analysis of colon LPLs. **b)** Percentages and numbers of total lymphocytes producing IFN-γ, GM-CSF, and IL-17A. Flow plots of CD4+ T-cells for IFN-γ production versus **c)** IL-10, **d)** GM-CSF and **e)** IL-17A. **f)** Gating of total CD4+ GM-CSF+ lymphocytes with corresponding flow plots of IFN-γ versus IL-17A. **g)** Gating of total GM-CSF+ lymphocytes with corresponding flow plots of IFN-γ versus IL-17A with bar plots of **h)** percentages (from live population) of lymphocytes producing GM-CSF alongside IL-17A, IFN-γ, or both. All flow analyses were performed on from *H. hepaticus* infected mice. **i)** Boxplots of log2(normalized read counts) of *Rorc* and *Foxp3* in bulk tissue LPL RNA-seq. **j)** Flow cytometry analysis of RORγt and Foxp3 transcription factor expression with **k)** percentages (top) and numbers (bottom) of CD4+ T cells producing of RORγt and/or Foxp3. **l)** Unsupervised hierarchical clustering of a pair-wise Spearman correlation of RNA-seq on sorted CD4+ T cells from colon LPL isolated from uninfected *Prdm1*^fl/fl^*Maf*^fl/fl^ and *Prdm1*^fl/fl^*Maf*^fl/fl^*Cd4^C^*^re^, and *H. hepaticus* infected *Prdm1*^fl/fl^*Maf*^fl/fl^ mice and mice with *Cd4*^Cre^-mediated deletion of either *Prdm1*, *Maf*, or both *Prdm1* and *Maf*. **m)** Number of differentially expressed genes identified in uninfected *Prdm1*^fl/fl^*Maf*^fl/fl^*Cd4^C^*^re^ and *H. hepaticus* infected mice compared to uninfected *Prdm1*^fl/fl^*Maf*^fl/fl^ controls (fold change >=1.5 and BH adjusted p value<0.05). **n)** Boxplots of log2(normalized read counts) of *Rorc* and *Foxp3* expression in sorted CD4+ from colon LPL. **o)** Unsupervised hierarchical clustering of a pair-wise Spearman correlation of ATAC-seq on sorted CD4+ T cells from colon LPL isolated from uninfected *Prdm1*^fl/fl^*Maf*^fl/fl^ and *Prdm1*^fl/fl^*Maf*^fl/fl^*Cd4^C^*^re^, and *H. hepaticus* infected *Prdm1*^fl/fl^*Maf*^fl/fl^ mice and mice with *Cd4*^Cre^-mediated deletion of either *Prdm1*, *Maf*, or both *Prdm1* and *Maf*. Data from n=3-5 mice.

**Extended Fig.5.**
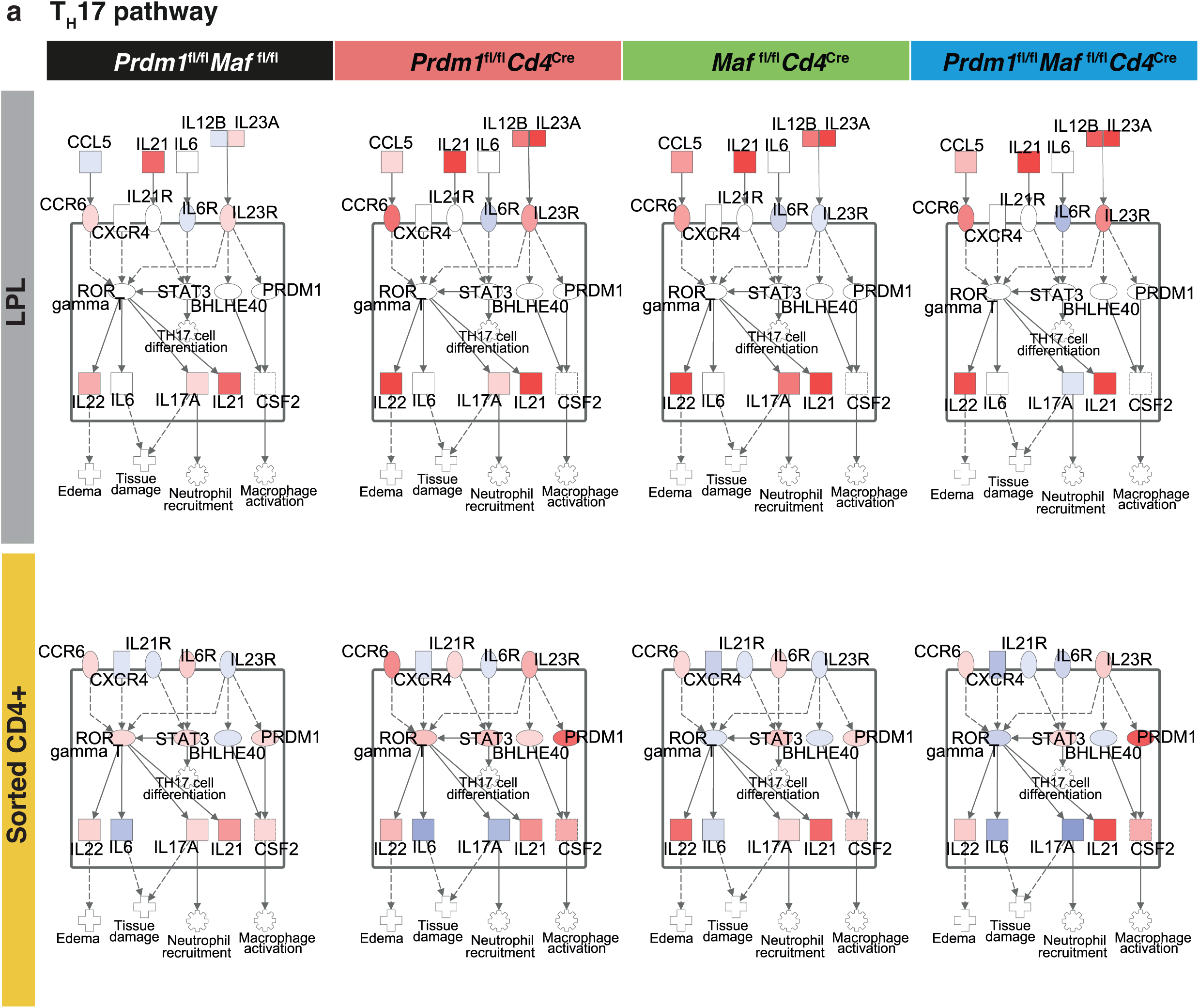
Bulk tissue colon LPL and sorted CD4+ T cell RNA-seq highlight upregulation of the Th17 pathway in *H.* hepaticus infected mice with *Cd4*^Cre^- mediated deletion of *Maf* when compared to infected *Prdm1*^fl/fl^*Maf*^fl/fl^ mice and mice with deletion of either *Prdm1* or both *Prdm1* and *Maf*. **a)** IPA was used to overlay bulk tissue LPL RNA-seq (top panel) and sorted CD4+ T cells RNA-seq (bottom panel) gene expression fold-changes in each *H. hepaticus* infected conditions relative to the uninfected *Prdm1*^fl/fl^*Maf*^fl/fl^ control into the “Th17 pathway”. A fixed scale of -5 (blue) to 3.5 (red) was kept between all conditions.

**Extended Fig.6.**
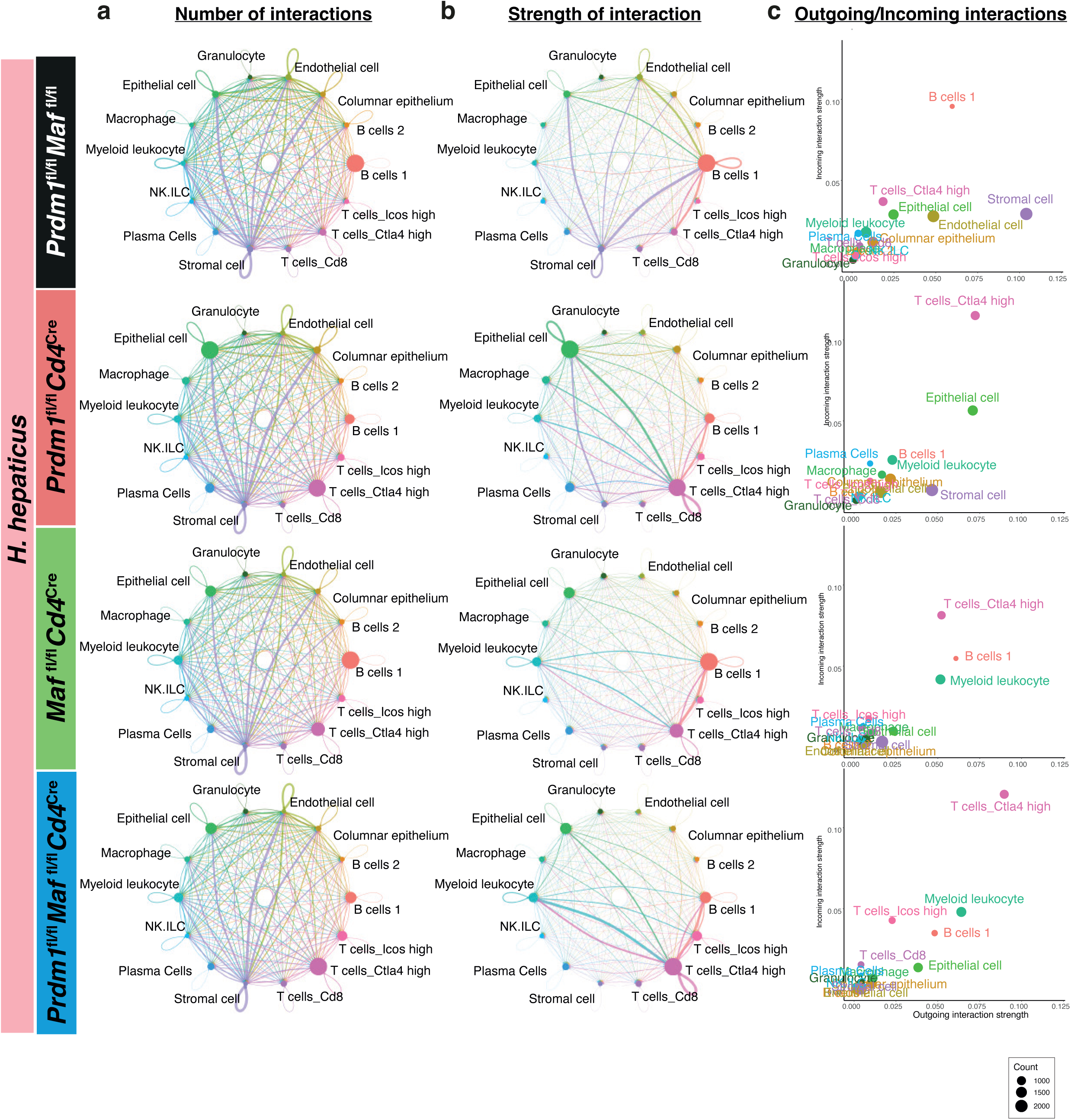
Inference of cell-to-cell communication networks in immune, stromal, epithelial and endothelial cell clusters from colonic LPL scRNA-seq. Cell-to-cell communication networks inferred using CellChat software from gene expression of ligands and their receptors in immune, stromal, epithelial and endothelial cell clusters from the colonic LPL scRNA-seq dataset. **a)** number and **b)** strength of interaction of cell-to-cell interactions, represented in the edge width, in and *H. hepaticus* infected *Prdm1*^fl/fl^*Maf*^fl/fl^ mice and mice with *Cd4*^Cre^-mediated deletion of either *Prdm1*, *Maf*, or both *Prdm1* and *Maf*. **c)** Plot of “Outgoing interaction strength“ against “Incoming interaction strength” across all *H. hepaticus* infected conditions. In **a-c)** node size is proportional to the number of cells in each experimental group, and the edges are colored based on the cell clusters expressing the outgoing signals.

**Extended Fig.7.**
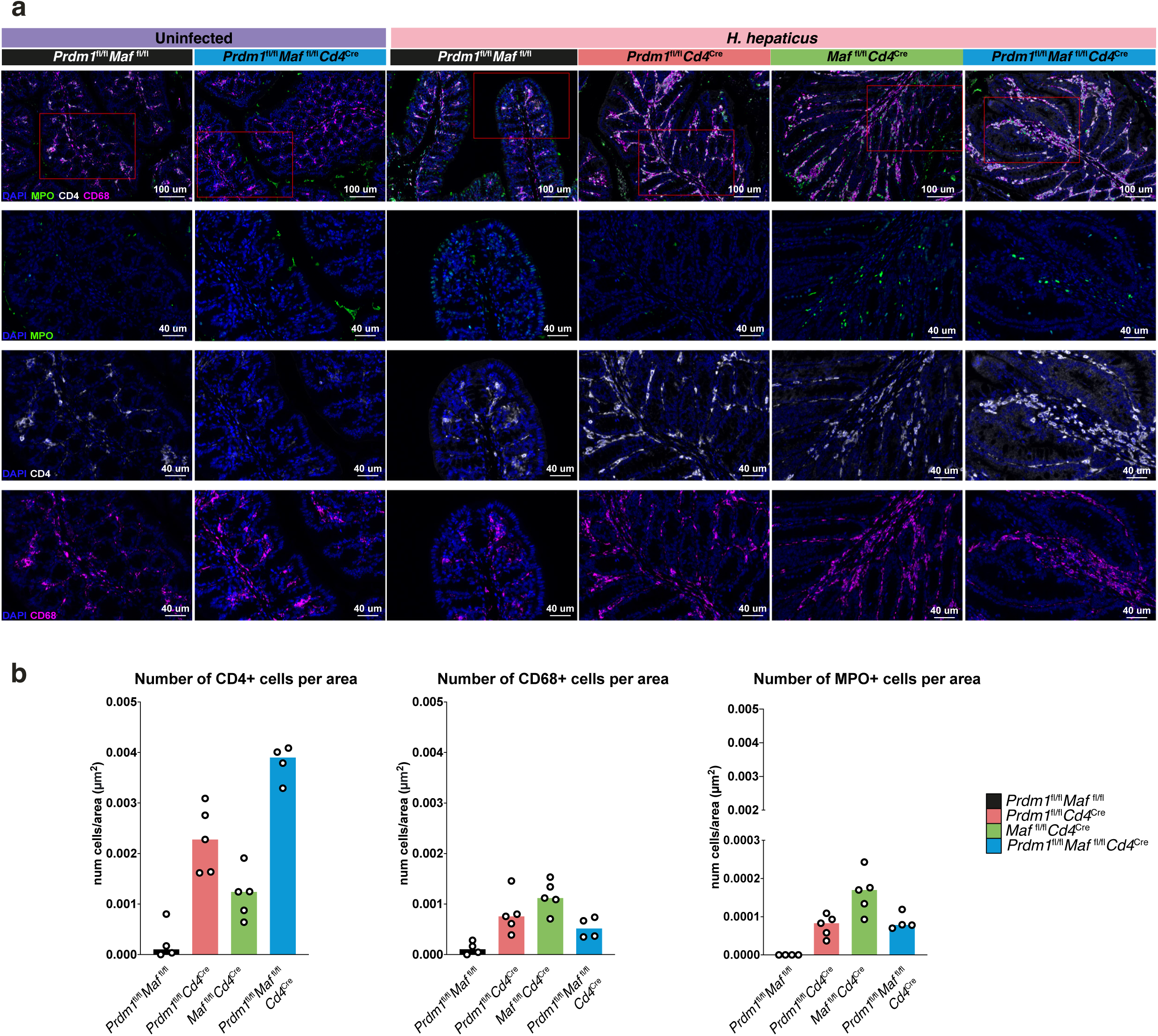
Immunofluorescence shows co-localization of neutrophils, CD4+ cells and mononuclear phagocytes in the colon. **a)** Representative images (n=4-5) of colon sections by immunofluorescence, staining for CD4+ T cells (CD4, white), mononuclear phagocytes (CD68, magenta), neutrophils (MPO, green) and nuclear staining (DAPI, blue) from uninfected *Prdm1*^fl/fl^*Maf*^fl/fl^ and *Prdm1*^fl/fl^*Maf*^fl/fl^*Cd4^C^*^re^, and *H. hepaticus* infected *Prdm1*^fl/fl^*Maf*^fl/fl^ mice and mice with *Cd4*^Cre^-mediated deletion of either *Prdm1*, *Maf*, or both *Prdm1* and *Maf*. Top row showing merged channels with a scale bar = 100μm; in the second row DAPI and MPO channels are shown with a scale bar = 40μm; in the third row DAPI and CD4 channels are shown with a scale bar = 40μm; and in the bottom row DAPI and CD68 channels are shown with a scale bar = 40μm. b) Bar graphs showing the median number of CD4+, CD68+ and MPO+ cells per area (μm^2^) in gut sections of each mouse from *H. hepaticus* infected *Prdm1*^fl/fl^*Maf*^fl/fl^ mice and mice with *Cd4*^Cre^-mediated deletion of either *Prdm1*, *Maf*, or both *Prdm1* and *Maf*. Data from n=4-5 mice.

**Extended Fig.8.**
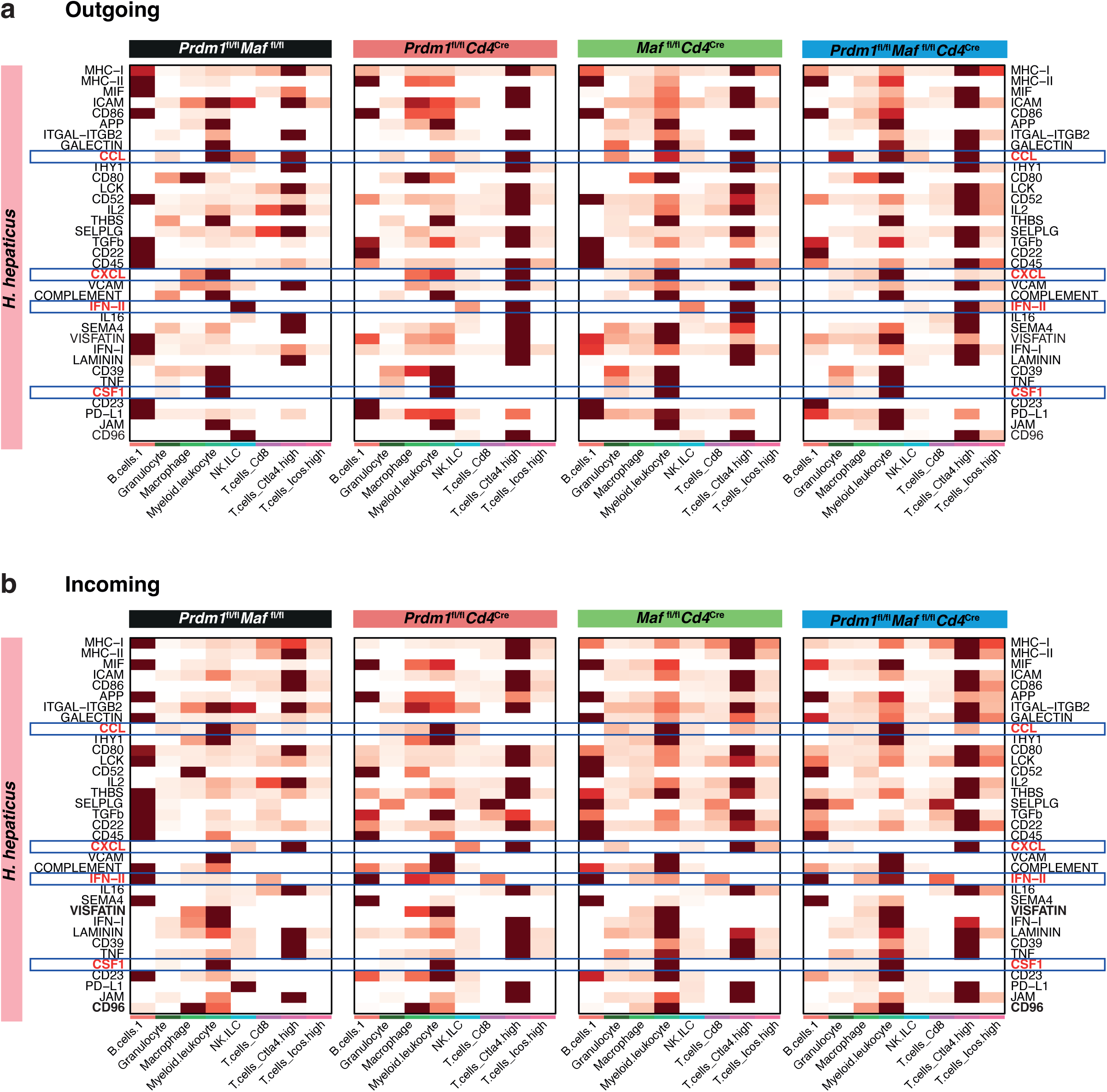
CellChat analysis of outgoing and incoming interaction strengths reveals a shift to increased interactions within immune and inflammation pathways in Tcells_Ctla4 high, Myeloid leukocytes and Macrophages in *H. hepaticus* infected mice with *Cd4*^Cre^-mediated deletion of *Prdm1*, *Maf,* or both *Prdm1* and *Maf* when compared to infected *Prdm1*^fl/fl^*Maf*^fl/fl^ mice. Heatmaps of **a)** outgoing and **b)** incoming interaction strengths from key pathways disrupted in the *H. hepaticus* infected *Prdm1*^fl/fl^*Maf*^fl/fl^*Cd4^C^*^re^ mice as compared to uninfected *Prdm1*^fl/fl^*Maf*^fl/fl^ control mice, interaction strength is shown per cell cluster and *H. hepaticus* infected condition.

**Extended Fig.9.**
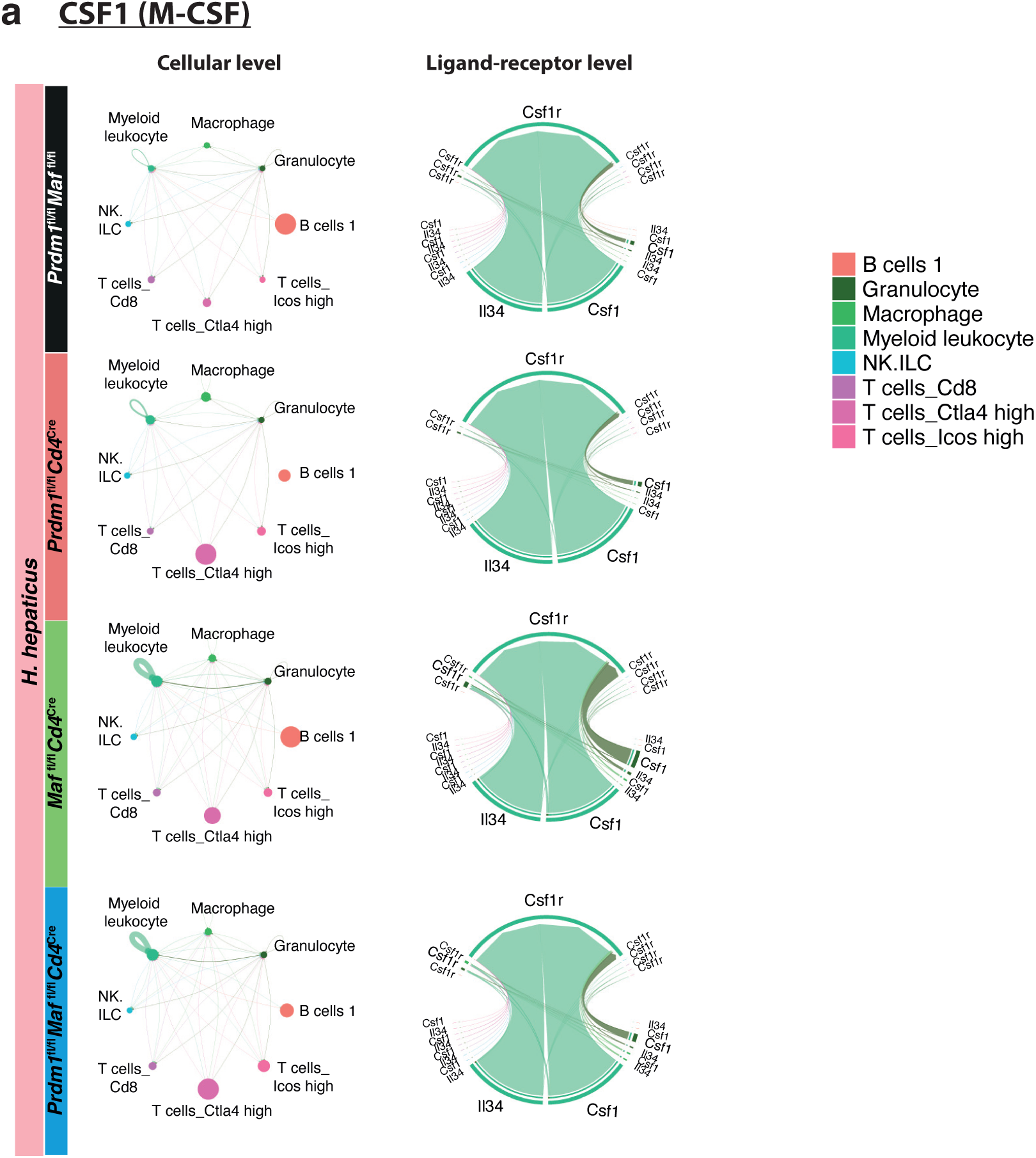
Putative cell-to-cell communication networks reveal an increase in Granulocyte and Myeloid leukocyte-driven M-CSF pathway in *H. hepaticus* infected mice with *Cd4*^Cre^-mediated deletion of *Maf* and both *Prdm1* and *Maf*. Cell-to-cell communication networks underlying the **a)** M-CSF pathway across all *H. hepaticus* infected *Prdm1*^fl/fl^*Maf*^fl/fl^ mice and mice with *Cd4*^Cre^-mediated deletion of either *Prdm1*, *Maf*, or both *Prdm1* and *Maf*. The chord plot has receiver cells at the top (incoming signaling) and transmitter cells (outgoing signaling) the bottom. The edges are colored based on the cell clusters expressing the outgoing signals.

